# Microorganism adhesion experimental study using silicon dioxide

**DOI:** 10.1101/388694

**Authors:** Roberts Lozins, Dzintars Ozoliņš

**Affiliations:** Faculty of Medicine, University of Latvia, Riga, Latvia

**Author notes:** Faculty of Medicine, University of Latvia, Riga, Latvia.

## Abstract

In this study, yeast, Gram positive and Gram negative bacteria were attached to silicon dioxide microparticles or silica in order to measure their absorbance, also known as physical absorption of light, changes using spectrophotometry. The goal of the study was to determine if spectrophotometry is an effective way to distinguish microorganisms and if microorganisms have an affinity for silicon dioxide since it is a suitable material for the production of prostheses. The experiment was done by examining the light absorption properties of yeast, Gram positive and Gram negative bacteria in a spectrophotometer with and without silicon dioxide microparticles. During the experiment there have been several promising results. First of all, the spectrophotometers presented graphs of yeast were noticeably different from the graphs of both Gram positive and Gram negative bacteria. Secondly, the absorption of light in both Gram positive and Gram negative bacteria at near infrared (700-1500 nm) wavelengths increased when silicon dioxide microparticles were added to the suspension, unlike yeast. When silicon dioxide microparticles were added to yeast, the absorption of light decreased during the whole wavelength interval of the spectrophotometer measurement. The results indicate that spectrophotometry could be used to distinguish yeast from bacteria and possibly bacteria from each other. The results also suggest that silicon dioxide should not be used in the production of prostheses since it could be a favourable material for the development of biofilms.

## Introductions

A key role in microorganism pathogenesis plays its adhesion to surfaces. Binding to the surface will give the microorganisms an opportunity to form biofilms, which are collection of microorganisms that are held together with a self-produced extracellular matrix. [1] Examining how the adhesion happens and what factor affect it would help to understand the pathogenesis of microorganisms better. Studies conducted by P. Loskill suggest that silica or silicon dioxide is a favourable surface for microorganism adhesion and it can be used to study the attracting forces between microorganisms and inorganic surfaces. [2]

To examine the adhesion of microorganisms to inorganic surfaces spectrophotometry was implied. A spectrophotometer is a well known device used for research and diagnostic purposes since 1940. It measures the transmission and absorption of light at certain wavelengths. [3] It can be used in medicine to determine the concentration of several substances in body fluids or the rate at which a clear substance changes its color in specific test, such as ELISA assay after adding an enzyme linked coloring agent. [4], [5] A spectrophotometer mechanism of action involves a light source, collimator, monochromator, wavelength selector and a photodetector. The collimator is a lens, that concentrates the beam of light to the monochromator. The monochromator works as a prism and it splits the beam of light into a large wavelength range. A wavelength selector selects a certain wavelength light beam, from the monochromators created wide range of wavelengths, which will interact with the examined substance. The photodetector will detect the light energy that passed through the examined substance. [6], [7]

There has been a modest amount of studys involving the adhesion properties of microorganisms involving spectrophotometry. One such study was conducted by A. Faltermeier, R. Bürgers and M. Rosentritt in 2008. The name of the study was “Bacterial adhesion of *Streptococcus mutans* to esthetic bracket materials”. In this study spectrophotometry was used to measure the quantity of biofilms formed by *Streptococcus mutans*. The study was aimed to determine if and how the materials used to make brackets affects the adherence of *Streptococcus mutants*. It was determined that the materials used to make brackets did not have a significant impact on the adhesion of *Streptococcus mutans* when compared one to the other. All plastic bracket materials had a similar bacterial colonization. [8]

Different study was conducted by D. Ozoliņš and A. Žilēviča and it was called “HIGHER SPEED OF YEAST ADHERENCE IN COMPARISON WITH SPEED OF BACTERIA ADHERENCE TO SILICA MICROPARTICLES”. The study was aimed to determine if there is a significant difference in the rate at which turbidity decreases or contrary adherence speed increases for bacteria and yeast with and without silicon dioxide microparticles. The method for taking measurement in the study was densitometry. It was determined that the decrease of turbidity when silicon dioxide was added was greater for both bacteria and yeast compared to bacteria and yeast without silicon dioxide. The study did also show that yeast binds quicker to silica than bacteria. [9] Other studies, such as “Surface Physicochemical Properties at the Micro and Nano Length Scales: Role on Bacterial Adhesion and *Xylella fastidiosa* Biofilm Development” by G. Lorite, R. Janissen and J. Clerici, indicate that silica is a more favourable surface for the development of biofilms than cellulose films, which are biotic surfaces [10].

A study conducted by B. Cousins called “Effects of a nanoparticulate silica substrate on cell attachment of *Candida albicans*”, indicated that silica nanoparticles reduced the growth and reproduction of *Candida albicans*. However, this study does not disprove the possibility that yeast binds quicker to silica than bacteria [11].

It is important to examine the adhesion of microorganisms since it plays a key role in there pathogenesis. It is also important to determine how microorganisms interact with silicon dioxide or silica, since it is commonly used in the production of different kinds of prostheses. [12], [13], [14]. Infection in a joint prosthesis may not be very common, but it is the most serious complication that can occur after implanting a prosthesis. The infection rate in the first 2 years after the implantation of a hip, shoulder or knee prostheses are below 2%, but the infection rate of an elbow prosthesis can reach up to 9%. [15]. It is also important to find new and faster ways to determine what microorganism is the cause of an infection.

The aim of the study was to examine if and what kind of microorganisms have an affinity for silicon dioxide and whether or not it is possible to distinguish one microorganism from another using spectrophotometry. The study included 4 yeast, 5 Gram positive and 8 Gram negative bacteria.

## Materials and methods

### Preparation of bacteria, yeast, silicon dioxide suspensions

The study was conducted in “The Microbiology laboratory of Traumatology and Orthopedic Hospital” in Latvia. To prepare a 4 McFarland unit silicon dioxide suspension, Sigma-Aldrich silicon dioxide microparticles with a mean particle size of 9-13 μm and density of 1.05 - 1.15 g/m were added to distilled water. Samples were taken from the suspension and placed in a test tube. The test tube was placed inside a densitometer and its turbidity was measured. The turbidity was measured using a DEN-1 densitometer. If needed, more silicon dioxide was added to distilled water till the densitometer showed the turbidity of 4±0,1 McFarland units in the sample test tube from the distilled water. Samples were taken from reference cultures of the microorganisms and placed in test tubes that were filled with 8 ml of distilled water. In total 17 microorganisms were examined and they were 5 reference cultures of Gram positive bacteria (*Enterococcus faecalis* ATCC (American Type Culture Collection) 29212, *Staphylococcus epidermidis* ATCC 12228, MSSA (Methicillin sensitive *Staphylococcus aureus*) ATCC 25923, MRSA (Methicillin resistant *Staphylococcus aureus*) ATCC 33591, *Bacillus spizizenii* ATCC 66338), 8 reference cultures of Gram negative bacteria *(Proteus mirabilis* ATCC 43071, *Enterobacter aerogenes* ATCC 13048, *Salmonella enteritidis* ATCC 13076, *Pseudomonas aeruginosa* ATCC 27853, *Citrobacter freundii* ATCC 438764, *Klebsiella pneumoniae* ATCC 700603, *Moraxella catarrhalis* ATCC 25238, *Escherichia coli* ATCC 25922) and 4 pure cultures of yeast: (*Candida kefyr, Candida glabrata, Candida krusei, Candida albicans* ATCC 10231). The created suspensions were placed inside the densitometer to measure there turbidity. If needed, more microorganism samples from the reference cultures were added till the densitometer showed the turbidity of 4±0,1 McFarland units.

### Filling cuvettes with microorganism and silicon dioxide suspensions

17 cuvettes were filled with syringes which contained 4 ml of the 4±0,1 McFarland unit suspensions of each microorganism. 17 other cuvettes were filled with syringes which contained 3.5 ml of the 4±0,1 McFarland unit suspensions of microorganisms. After that 0.5 ml of 4±0,1 McFarland unit silicon dioxide suspension was added to these cuvettes. 1 cuvette was filled with 4 ml of distilled water and another one was filled with 4 ml of 4±0,1 McFarland unit silicon dioxide suspension.

### Calibration of the spectrophotometer and measuring the absorbance of microorganisms

The spectrophoteometers “Mode” was set to “Spectrum”. A cuvette with distilled water was placed in the spectrophotometer and the “Base Core” option was selected. This made sure that the light that went trough distilled water in all wavelengths measured by the spectrophotometer was 100% of light detected by the photodetector or 0 absorbance units. The spectrophotometer was set to measure the absorbance in a wavelength interval of 285-1100 nm. The speed of the measurement was set to “Fast” and the distance between each measured wavelength was set to 1 nm. Cuvette with the measured microorganism was placed in the spectrophotometer. “Start” button was pressed to start the measurement. The measurements were conducted again with the wavelength interval of 285-700 nm. The difference of each measured wavelength was set to 1 nm.

### Comparing microorganisms

All results were converted in the spectrophotometer from its used SPC format to CSV format. The results were sent to a flash drive and placed on a computer. Each microorganisms measurement contained the wavelengths of the measurement and the absorbance units at the current wavelength. All measured absorbances of microorganisms with the wavelength interval of 285-1100 nm and without silicon dioxide microparticles were compared to each other using t-test: Two-Sample assuming unequal variances. “Alpha” was set to 0.05 and “Hypothesized Mean Difference” was set to 0. The comparisons were done in *MS Excel* 2016. Same comparisons were done for microorganism measurements at 285-1100 nm with silicon dioxide microparticles, 285-700 nm without silicon dioxide microparticles and at 285-700 nm with silicon dioxide microparticles. A significant difference was assumed if the *p* value was <0.05. Acquired results were gathered in tables in *MS Office* 2016 and graphs in *MS Excell*.

## Results

### Microorganisms comparison at 285-1100 nm

Examining the results presented in Table 1, which are measurement comparisons at 285-1100 nm without silicon dioxide, in 99/136 (72.8%) comparisons it was possible to distinguish one microorganism from another, assuming *p* value <0.05. In 52/52 (100%) comparisons in which yeast were compared to bacteria there was a significant difference. In 41/78 (52.6%) comparisons in which bacteria were compared to each other there was a significant difference. In 5/6 (83.3%) comparisons that were only between yeast there was a significant difference.

**Table 1.**
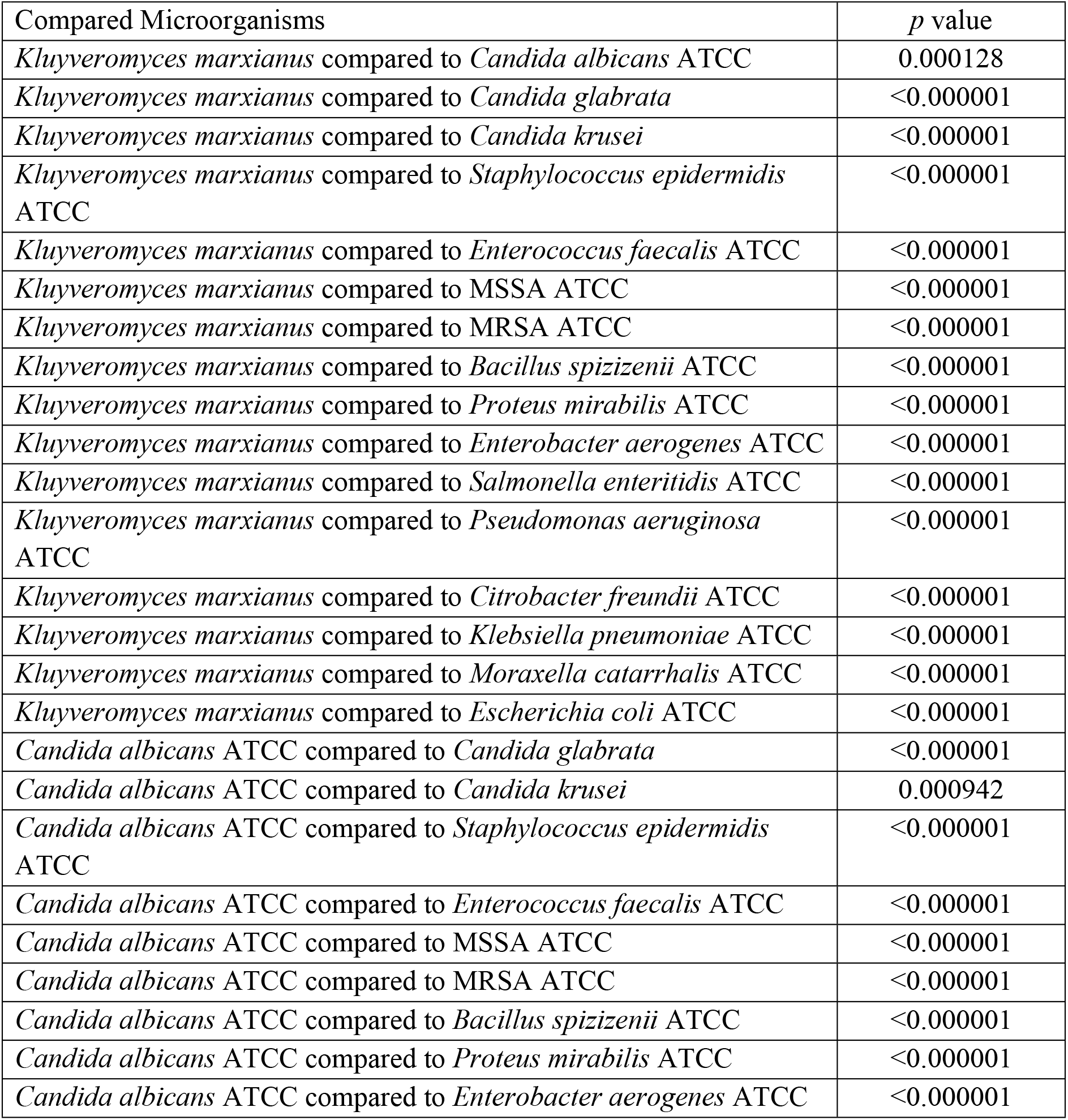
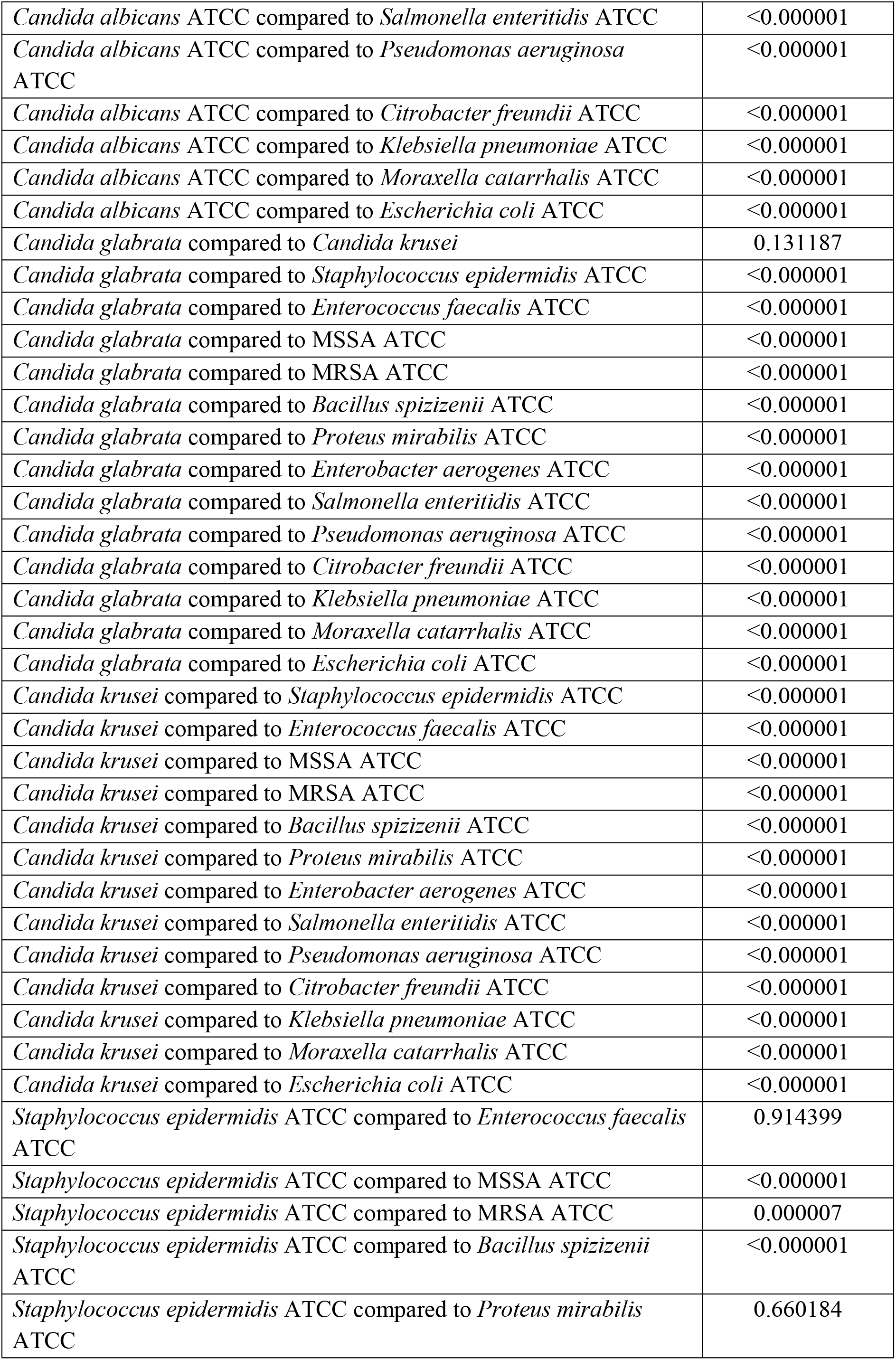
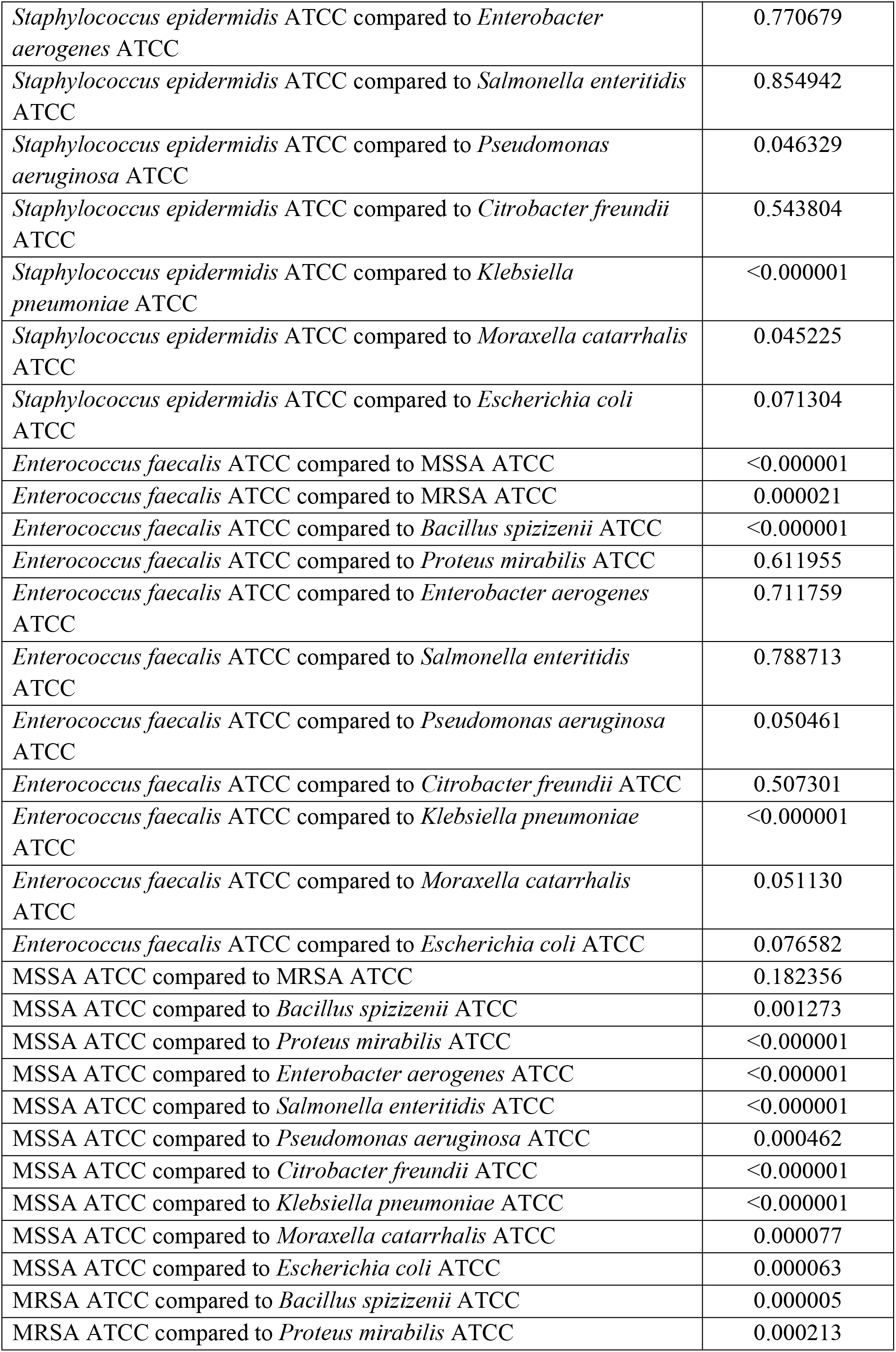
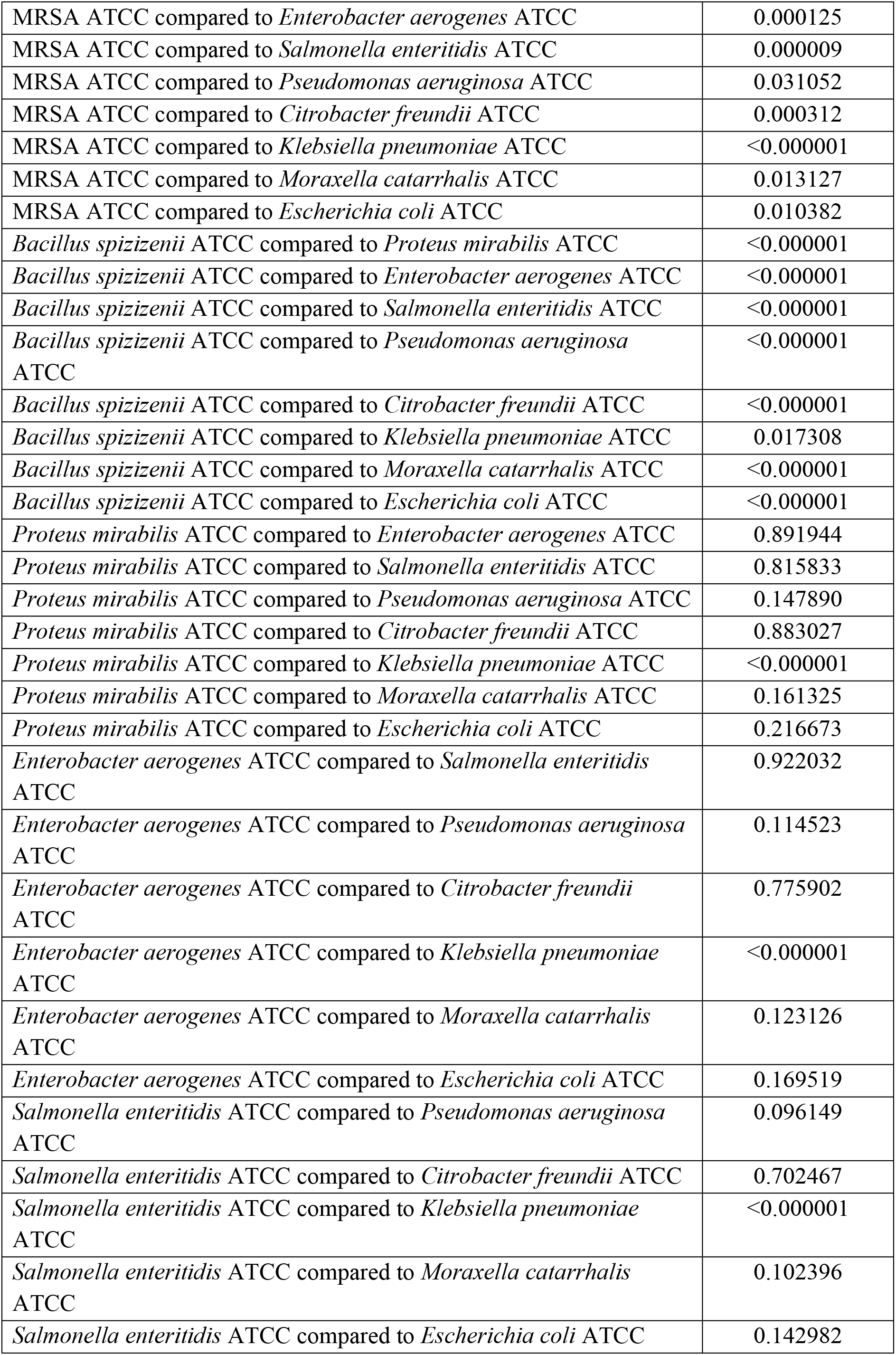
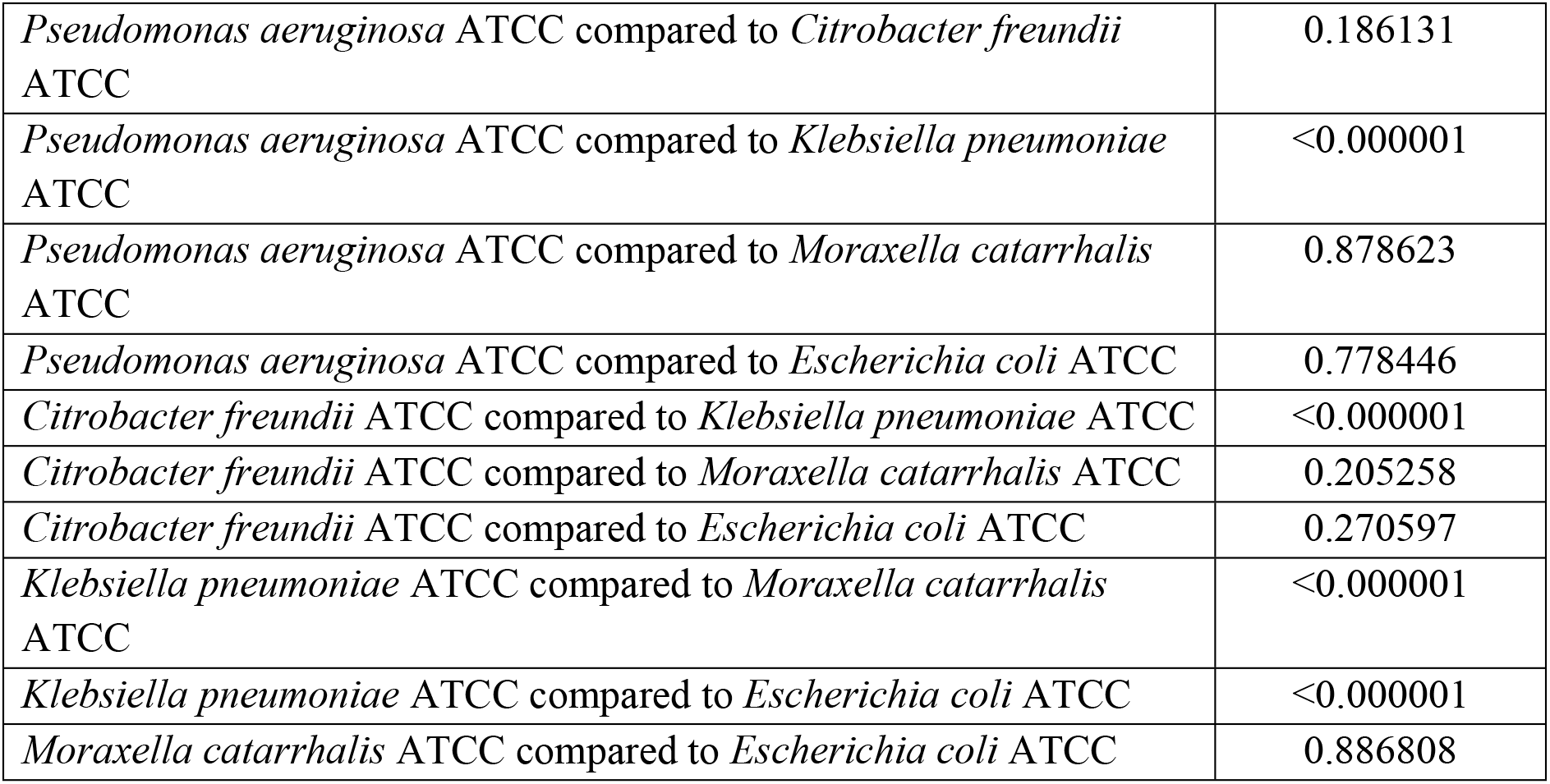
Comparison of microorganisms at 285-1100 nm without silicon dioxide using t-Test: Two-Sample Assuming Unequal Variances.

Examining the results presented in Table 2, which are measurement comparisons at 285-1100 nm with silicon dioxide, in 109/136 (81.1%) comparisons it was possible to distinguish one microorganism from another. In 52/52 (100%) comparisons in which yeast were compared to bacteria there was a significant difference. In 54/78 (69.2%) comparisons in which bacteria were compared to each other there was a significant difference. In 3/6 (50%) comparisons that were only between yeast there was a significant difference.

**Table 2.** Comparison of microorganisms at 285-1100 nm with silicon dioxide using t-Test: Two-Sample Assuming Unequal Variances.

| Compared Microorganisms | *p* value |
| --- | --- |
| *Kluyveromyces marxianus* compared to *Candida albicans* ATCC | 0.006617 |
| *Kluyveromyces marxianus* compared to *Candida glabrata* | 0.278350 |
| *Kluyveromyces marxianus* compared to *Candida krusei* | 0.014212 |
| *Kluyveromyces marxianus* compared to *Staphylococcus epidermidis* ATCC | <0.000001 |
| *Kluyveromyces marxianus* compared to *Enterococcus faecalis* ATCC | <0.000001 |
| *Kluyveromyces marxianus* compared to MSSA ATCC | <0.000001 |
| *Kluyveromyces marxianus* compared to MRSA ATCC | <0.000001 |
| *Kluyveromyces marxianus* compared to *Bacillus spizizenii* ATCC | <0.000001 |
| *Kluyveromyces marxianus* compared to *Proteus mirabilis* ATCC | <0.000001 |
| *Kluyveromyces marxianus* compared to *Enterobacter aerogenes* ATCC | <0.000001 |
| *Kluyveromyces marxianus* compared to *Salmonella enteritidis* ATCC | <0.000001 |
| *Kluyveromyces marxianus* compared to *Pseudomonas aeruginosa* ATCC | <0.000001 |
| *Kluyveromyces marxianus* compared to *Citrobacter freundii* ATCC | <0.000001 |
| *Kluyveromyces marxianus* compared to *Klebsiella pneumoniae* ATCC | <0.000001 |
| *Kluyveromyces marxianus* compared to *Moraxella catarrhalis* ATCC | <0.000001 |
| *Kluyveromyces marxianus* compared to *Escherichia coli* ATCC | <0.000001 |
| *Candida albicans* ATCC compared to *Candida glabrata* | <0.000001 |
| *Candida albicans* ATCC compared to *Candida krusei* | 0.889397 |
| *Candida albicans* ATCC compared to *Staphylococcus epidermidis* ATCC | <0.000001 |
| *Candida albicans* ATCC compared to *Enterococcus faecalis* ATCC | <0.000001 |
| *Candida albicans* ATCC compared to MSSA ATCC | <0.000001 |
| *Candida albicans* ATCC compared to MRSA ATCC | <0.000001 |
| *Candida albicans* ATCC compared to *Bacillus spizizenii* ATCC | <0.000001 |
| *Candida albicans* ATCC compared to *Proteus mirabilis* ATCC | <0.000001 |
| *Candida albicans* ATCC compared to *Enterobacter aerogenes* ATCC | <0.000001 |
| *Candida albicans* ATCC compared to *Salmonella enteritidis* ATCC | <0.000001 |
| *Candida albicans* ATCC compared to *Pseudomonas aeruginosa* ATCC | <0.000001 |
| *Candida albicans* ATCC compared to *Citrobacter freundii* ATCC | <0.000001 |
| *Candida albicans* ATCC compared to *Klebsiella pneumoniae* ATCC | <0.000001 |
| *Candida albicans* ATCC compared to *Moraxella catarrhalis* ATCC | <0.000001 |
| *Candida albicans* ATCC compared to *Escherichia coli* ATCC | <0.000001 |
| *Candida glabrata* compared to *Candida krusei* | 0.000133 |
| *Candida glabrata* compared to *Staphylococcus epidermidis* ATCC | <0.000001 |
| *Candida glabrata* compared to *Enterococcus faecalis* ATCC | <0.000001 |
| *Candida glabrata* compared to MSSA ATCC | <0.000001 |
| *Candida glabrata* compared to MRSA ATCC | <0.000001 |
| *Candida glabrata* compared to *Bacillus spizizenii* ATCC | <0.000001 |
| *Candida glabrata* compared to *Proteus mirabilis* ATCC | <0.000001 |
| *Candida glabrata* compared to *Enterobacter aerogenes* ATCC | <0.000001 |
| *Candida glabrata* compared to *Salmonella enteritidis* ATCC | <0.000001 |
| *Candida glabrata* compared to *Pseudomonas aeruginosa* ATCC | <0.000001 |
| *Candida glabrata* compared to *Citrobacter freundii* ATCC | <0.000001 |
| *Candida glabrata* compared to *Klebsiella pneumoniae* ATCC | <0.000001 |
| *Candida glabrata* compared to *Moraxella catarrhalis* ATCC | <0.000001 |
| *Candida glabrata* compared to *Escherichia coli* ATCC | <0.000001 |
| *Candida krusei* compared to *Staphylococcus epidermidis* ATCC | <0.000001 |
| *Candida krusei* compared to *Enterococcus faecalis* ATCC | <0.000001 |
| *Candida krusei* compared to MSSA ATCC | <0.000001 |
| *Candida krusei* compared to MRSA ATCC | <0.000001 |
| *Candida krusei* compared to *Bacillus spizizenii* ATCC | <0.000001 |
| *Candida krusei* compared to *Proteus mirabilis* ATCC | <0.000001 |
| *Candida krusei* compared to *Enterobacter aerogenes* ATCC | <0.000001 |
| *Candida krusei* compared to *Salmonella enteritidis* ATCC | <0.000001 |
| *Candida krusei* compared to *Pseudomonas aeruginosa* ATCC | <0.000001 |
| *Candida krusei* compared to *Citrobacter freundii* ATCC | <0.000001 |
| *Candida krusei* compared to *Klebsiella pneumoniae* ATCC | <0.000001 |
| *Candida krusei* compared to *Moraxella catarrhalis* ATCC | <0.000001 |
| *Candida krusei* compared to *Escherichia coli* ATCC | <0.000001 |
| *Staphylococcus epidermidis* ATCC compared to *Enterococcus faecalis* ATCC | 0.037622 |
| *Staphylococcus epidermidis* ATCC compared to MSSA ATCC | 0.000184 |
| *Staphylococcus epidermidis* ATCC compared to MRSA ATCC | 0.722643 |
| *Staphylococcus epidermidis* ATCC compared to *Bacillus spizizenii* ATCC | <0.000001 |
| *Staphylococcus epidermidis* ATCC compared to *Proteus mirabilis* ATCC | 0.007244 |
| *Staphylococcus epidermidis* ATCC compared to *Enterobacter aerogenes* ATCC | 0.000257 |
| *Staphylococcus epidermidis* ATCC compared to *Salmonella enteritidis* ATCC | 0.000004 |
| *Staphylococcus epidermidis* ATCC compared to *Pseudomonas aeruginosa* ATCC | 0.140340 |
| *Staphylococcus epidermidis* ATCC compared to *Citrobacter freundii* ATCC | 0.000026 |
| *Staphylococcus epidermidis* ATCC compared to *Klebsiella pneumoniae* ATCC | 0.00552 |
| *Staphylococcus epidermidis* ATCC compared to *Moraxella catarrhalis* ATCC | 0.071354 |
| *Staphylococcus epidermidis* ATCC compared to *Escherichia coli* ATCC | 0.654009 |
| *Enterococcus faecalis* ATCC compared to MSSA ATCC | <0.000001 |
| *Enterococcus faecalis* ATCC compared to MRSA ATCC | 0.027201 |
| *Enterococcus faecalis* ATCC compared to *Bacillus spizizenii* ATCC | <0.000001 |
| *Enterococcus faecalis* ATCC compared to *Proteus mirabilis* ATCC | 0.535533 |
| *Enterococcus faecalis* ATCC compared to *Enterobacter aerogenes* ATCC | 0.116586 |
| *Enterococcus faecalis* ATCC compared to *Salmonella enteritidis* ATCC | 0.012177 |
| *Enterococcus faecalis* ATCC compared to *Pseudomonas aeruginosa* ATCC | 0.649447 |
| *Enterococcus faecalis* ATCC compared to *Citrobacter freundii* ATCC | 0.043448 |
| *Enterococcus faecalis* ATCC compared to *Klebsiella pneumoniae* ATCC | 0.000002 |
| *Enterococcus faecalis* ATCC compared to *Moraxella catarrhalis* ATCC | 0.772104 |
| *Enterococcus faecalis* ATCC compared to *Escherichia coli* ATCC | 0.131859 |
| MSSA ATCC compared to MRSA ATCC | 0.002679 |
| MSSA ATCC compared to *Bacillus spizizenii* ATCC | 0.001238 |
| MSSA ATCC compared to *Proteus mirabilis* ATCC | <0.000001 |
| MSSA ATCC compared to *Enterobacter aerogenes* ATCC | <0.000001 |
| MSSA ATCC compared to *Salmonella enteritidis* ATCC | <0.000001 |
| MSSA ATCC compared to *Pseudomonas aeruginosa* ATCC | 0.000002 |
| MSSA ATCC compared to *Citrobacter freundii* ATCC | <0.000001 |
| MSSA ATCC compared to *Klebsiella pneumoniae* ATCC | 0.237971 |
| MSSA ATCC compared to *Moraxella catarrhalis* ATCC | <0.000001 |
| MSSA ATCC compared to *Escherichia coli* ATCC | 0.000103 |
| MRSA ATCC compared to *Bacillus spizizenii* ATCC | <0.000001 |
| MRSA ATCC compared to *Proteus mirabilis* ATCC | 0.005793 |
| MRSA ATCC compared to *Enterobacter aerogenes* ATCC | 0.000271 |
| MRSA ATCC compared to *Salmonella enteritidis* ATCC | 0.000006 |
| MRSA ATCC compared to *Pseudomonas aeruginosa* ATCC | 0.096462 |
| MRSA ATCC compared to *Citrobacter freundii* ATCC | 0.000036 |
| MRSA ATCC compared to *Klebsiella pneumoniae* ATCC | 0.036991 |
| MRSA ATCC compared to *Moraxella catarrhalis* ATCC | 0.050037 |
| MRSA ATCC compared to *Escherichia coli* ATCC | 0.461519 |
| *Bacillus spizizenii* ATCC compared to *Proteus mirabilis* ATCC | <0.000001 |
| *Bacillus spizizenii* ATCC compared to *Enterobacter aerogenes* ATCC | <0.000001 |
| *Bacillus spizizenii* ATCC compared to *Salmonella enteritidis* ATCC | <0.000001 |
| *Bacillus spizizenii* ATCC compared to *Pseudomonas aeruginosa* ATCC | <0.000001 |
| *Bacillus spizizenii* ATCC compared to *Citrobacter freundii* ATCC | <0.000001 |
| *Bacillus spizizenii* ATCC compared to *Klebsiella pneumoniae* ATCC | 0.000001 |
| *Bacillus spizizenii* ATCC compared to *Moraxella catarrhalis* ATCC | <0.000001 |
| *Bacillus spizizenii* ATCC compared to *Escherichia coli* ATCC | <0.000001 |
| *Proteus mirabilis* ATCC compared to *Enterobacter aerogenes* ATCC | 0.347364 |
| *Proteus mirabilis* ATCC compared to *Salmonella enteritidis* ATCC | 0.061169 |
| *Proteus mirabilis* ATCC compared to *Pseudomonas aeruginosa* ATCC | 0.301413 |
| *Proteus mirabilis* ATCC compared to *Citrobacter freundii* ATCC | 0.173284 |
| *Proteus mirabilis* ATCC compared to *Klebsiella pneumoniae* ATCC | <0.000001 |
| *Proteus mirabilis* ATCC compared to *Moraxella catarrhalis* ATCC | 0.362275 |
| *Proteus mirabilis* ATCC compared to *Escherichia coli* ATCC | 0.036913 |
| *Enterobacter aerogenes* ATCC compared to *Salmonella enteritidis* ATCC | 0.348114 |
| *Enterobacter aerogenes* ATCC compared to *Pseudomonas aeruginosa* ATCC | 0.053585 |
| *Enterobacter aerogenes* ATCC compared to *Citrobacter freundii* ATCC | 0.696695 |
| *Enterobacter aerogenes* ATCC compared to *Klebsiella pneumoniae* ATCC | <0.000001 |
| *Enterobacter aerogenes* ATCC compared to *Moraxella catarrhalis* ATCC | 0.061709 |
| *Enterobacter aerogenes* ATCC compared to *Escherichia coli* ATCC | 0.002674 |
| *Salmonella enteritidis* ATCC compared to *Pseudomonas aeruginosa* ATCC | 0.004847 |
| *Salmonella enteritidis* ATCC compared to *Citrobacter freundii* ATCC | 0.561159 |
| *Salmonella enteritidis* ATCC compared to *Klebsiella pneumoniae* ATCC | <0.000001 |
| *Salmonella enteritidis* ATCC compared to *Moraxella catarrhalis* ATCC | 0.004938 |
| *Salmonella enteritidis* ATCC compared to *Escherichia coli* ATCC | 0.000095 |
| *Pseudomonas aeruginosa* ATCC compared to *Citrobacter freundii* ATCC | <0.000001 |
| *Pseudomonas aeruginosa* ATCC compared to *Klebsiella pneumoniae* ATCC | <0.000001 |
| *Pseudomonas aeruginosa* ATCC compared to *Moraxella catarrhalis* ATCC | 0.852092 |
| *Pseudomonas aeruginosa* ATCC compared to *Escherichia coli* ATCC | 0.327486 |
| *Citrobacter freundii* ATCC compared to *Klebsiella pneumoniae* ATCC | <0.000001 |
| *Citrobacter freundii* ATCC compared to *Moraxella catarrhalis* ATCC | 0.019583 |
| *Citrobacter freundii* ATCC compared to *Escherichia coli* ATCC | 0.000489 |
| *Klebsiella pneumoniae* ATCC compared to *Moraxella catarrhalis* ATCC | 0.000008 |
| *Klebsiella pneumoniae* ATCC compared to *Escherichia coli* ATCC | 0.002762 |
| *Moraxella catarrhalis* ATCC compared to *Escherichia coli* ATCC | <0.000001 |

Examining the results presented in Table 3, which are measurement comparisons at 285-700 nm without silicon dioxide, in 106/136 (77.9%) comparisons it was possible to distinguish one microorganism from another. In 44/52 (84.6%) comparisons in which yeast were compared to bacteria there was a significant difference. In 56/78 (71.8%) comparisons in which bacteria were compared to each other there was a significant difference. In 6/6 (100%) comparisons that were only between yeast there was a significant difference.

**Table 3.**
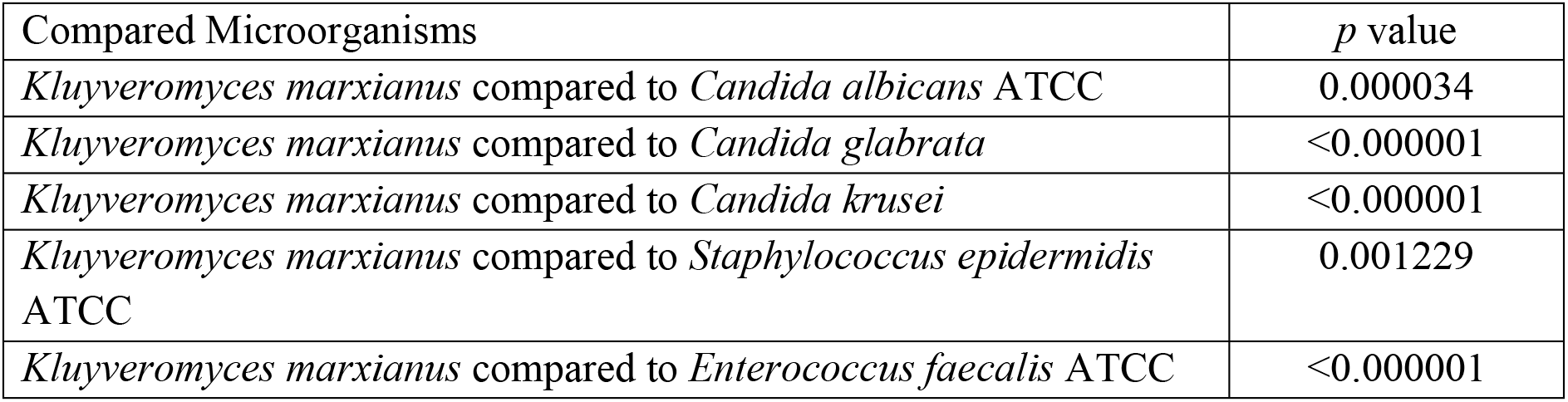
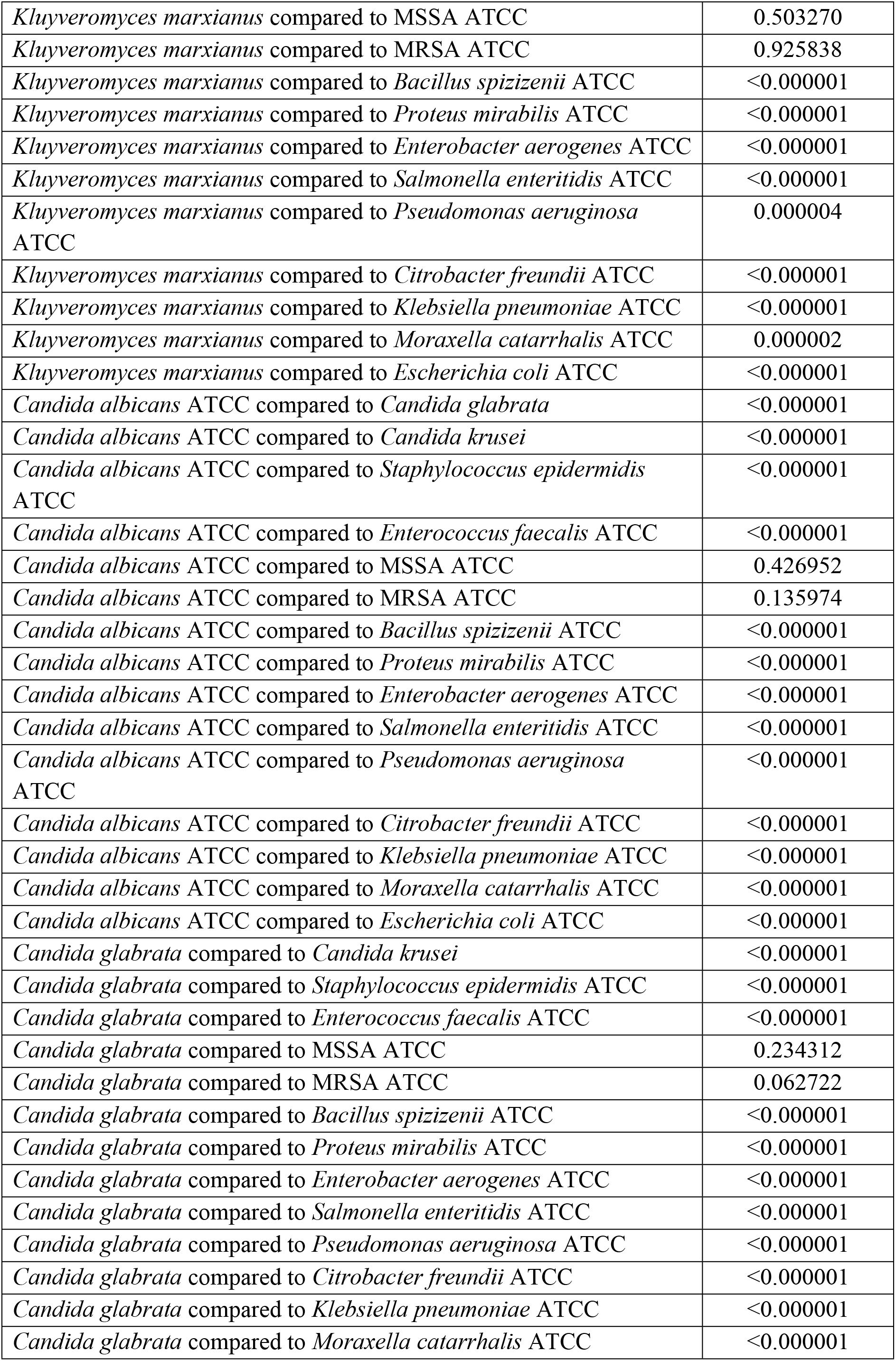
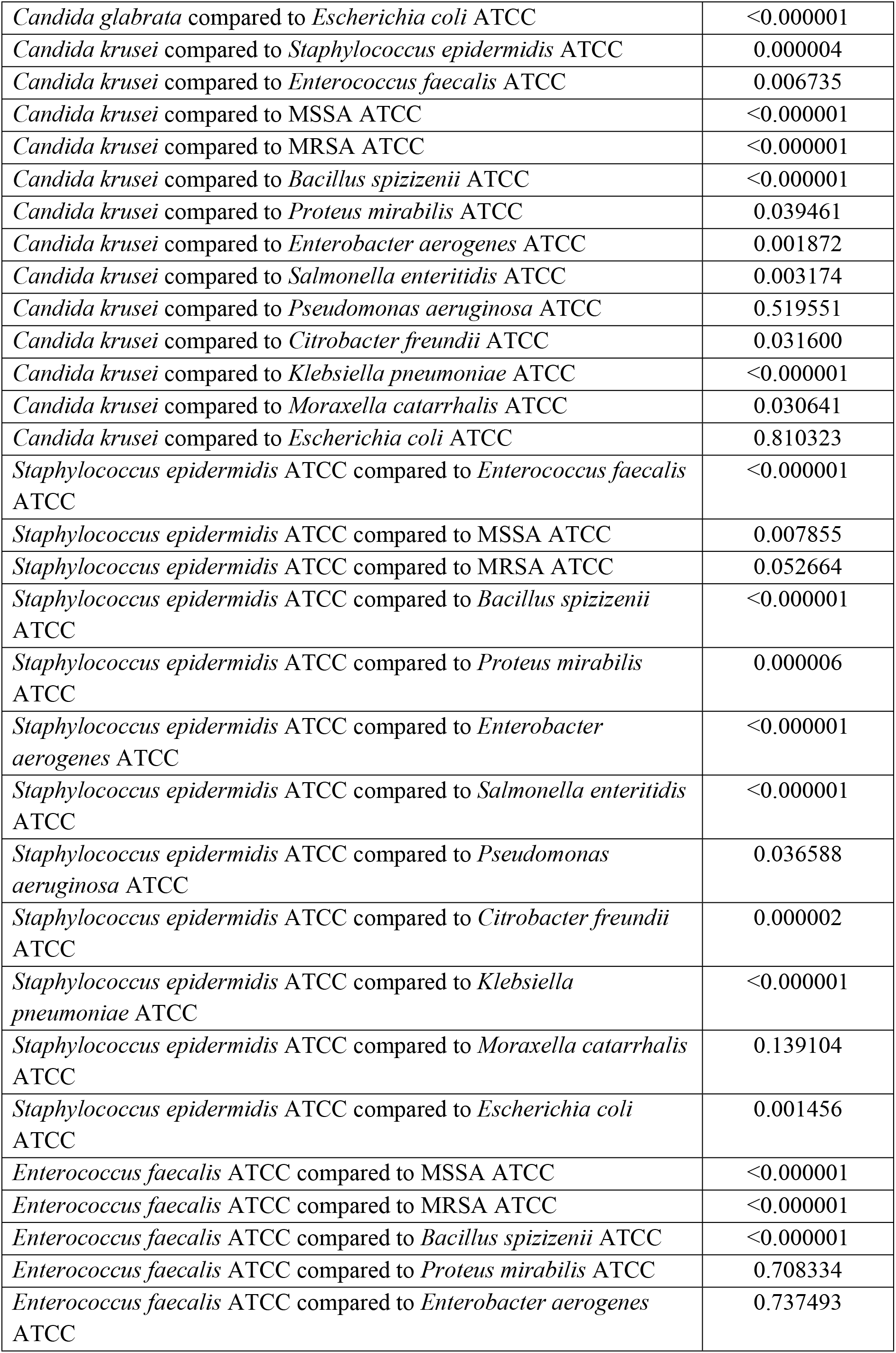
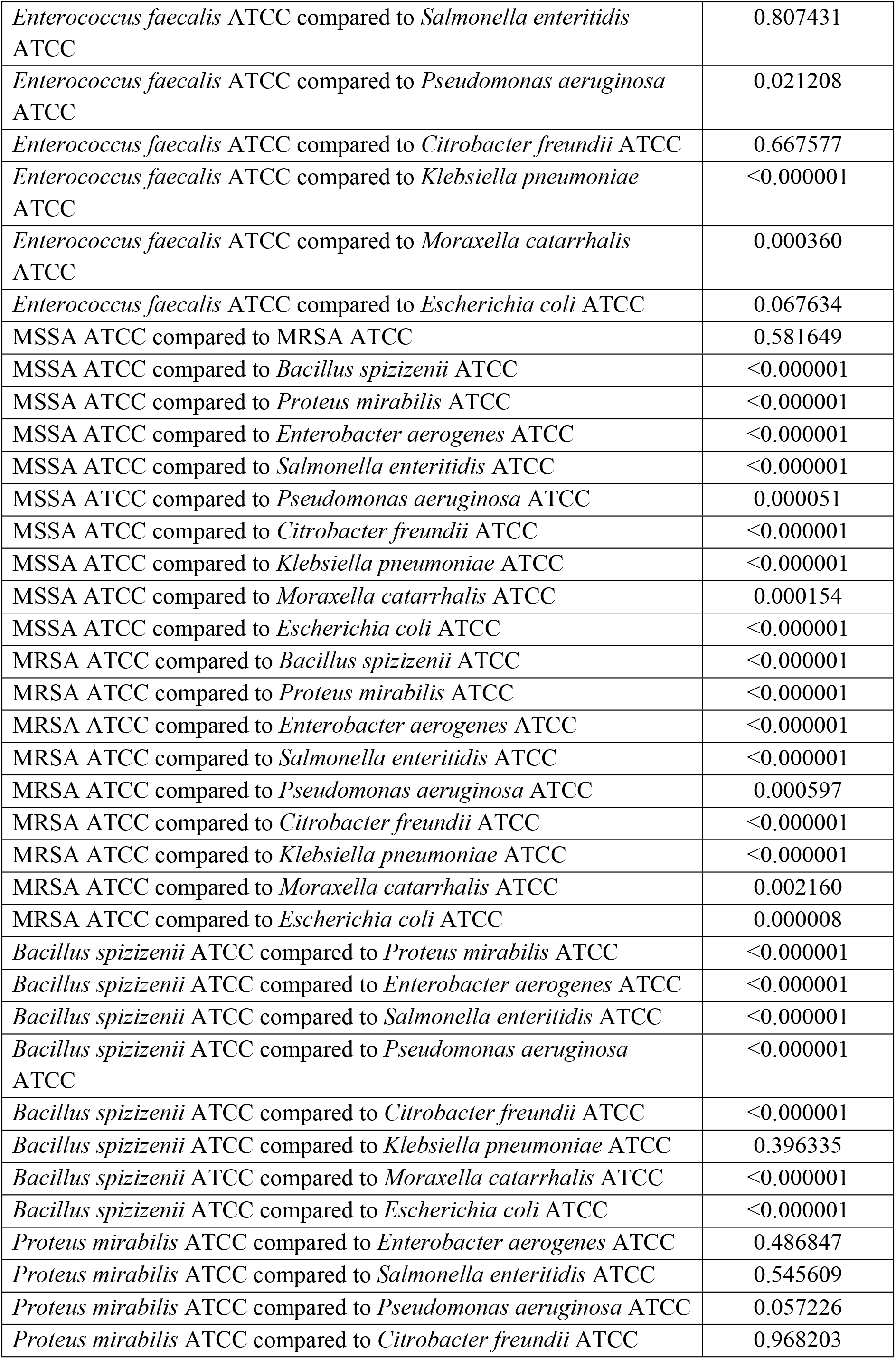
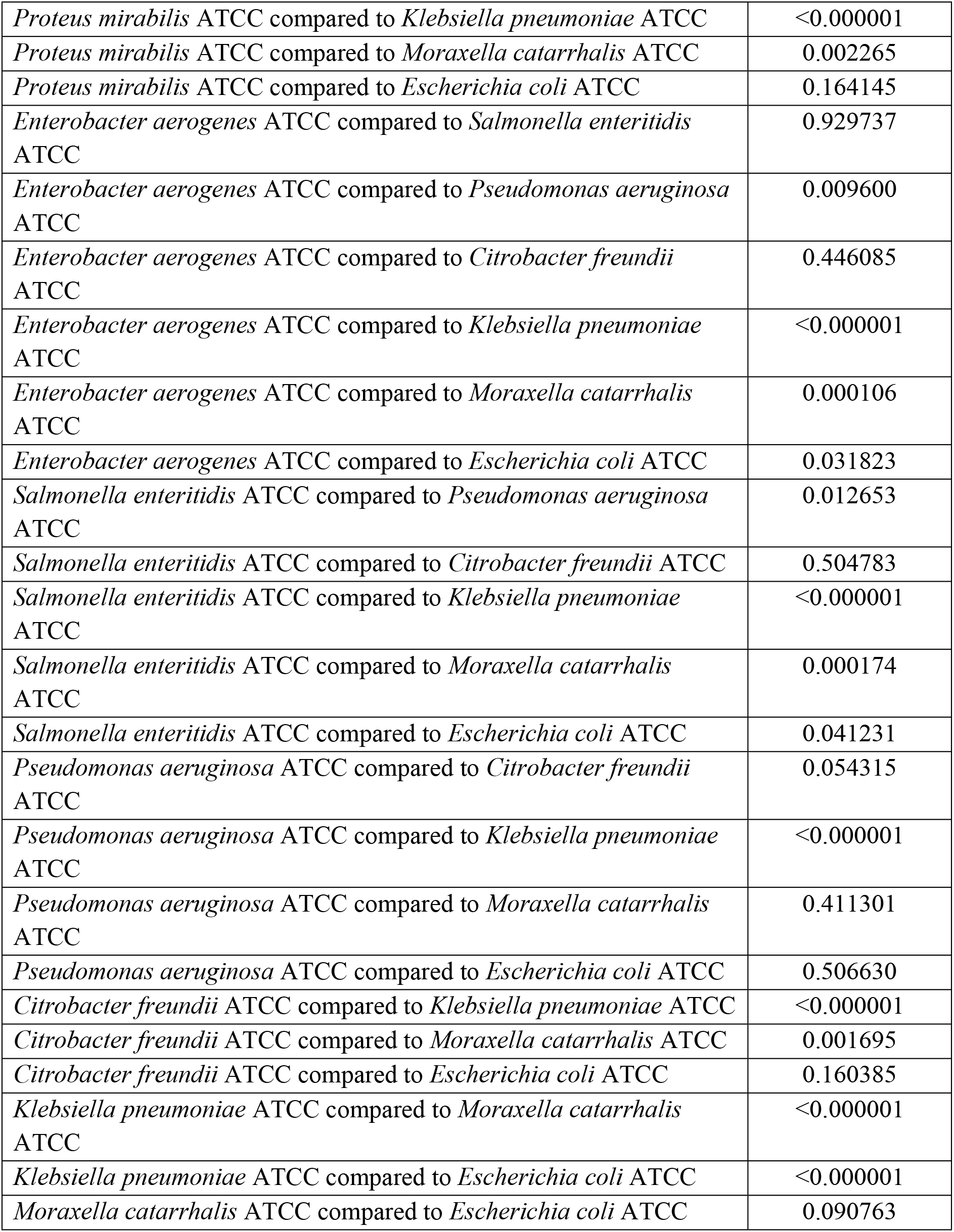
Comparison of microorganisms at 285-700 nm without silicon dioxide using t-Test: Two-Sample Assuming Unequal Variances.

### Microorganisms comparison at 285-700 nm

Examining the results presented in Table 4, which are measurement comparisons at 285-700 nm with silicon dioxide, in 107/136 (78.7%) comparisons it was possible to distinguish one microorganism from another. In 44/52 (84.6%) comparisons in which yeast were compared to bacteria there was a significant difference. In 58/78 (74.6%) comparisons in which bacteria were compared to each other there was a significant difference. In 5/6 (83.3%) comparisons that were only between yeast there was a significant difference.

**Table 4.** Comparison of microorganisms at 285-700 nm with silicon dioxide using t-Test: Two-Sample Assuming Unequal Variances.

| Compared Microorganisms | *p* value |
| --- | --- |
| *Kluyveromyces marxianus* compared to *Candida albicans* ATCC | 0.601437 |
| *Kluyveromyces marxianus* compared to *Candida glabrata* | <0.000001 |
| *Kluyveromyces marxianus* compared to *Candida krusei* | <0.000001 |
| *Kluyveromyces marxianus* compared to *Staphylococcus epidermidis* ATCC | 0.993349 |
| *Kluyveromyces marxianus* compared to *Enterococcus faecalis* ATCC | <0.000001 |
| *Kluyveromyces marxianus* compared to MSSA ATCC | 0.518039 |
| *Kluyveromyces marxianus* compared to MRSA ATCC | <0.000001 |
| *Kluyveromyces marxianus* compared to *Bacillus spizizenii* ATCC | <0.000001 |
| *Kluyveromyces marxianus* compared to *Proteus mirabilis* ATCC | <0.000001 |
| *Kluyveromyces marxianus* compared to *Enterobacter aerogenes* ATCC | <0.000001 |
| *Kluyveromyces marxianus* compared to *Salmonella enteritidis* ATCC | <0.000001 |
| *Kluyveromyces marxianus* compared to *Pseudomonas aeruginosa* ATCC | <0.000001 |
| *Kluyveromyces marxianus* compared to *Citrobacter freundii* ATCC | <0.000001 |
| *Kluyveromyces marxianus* compared to *Klebsiella pneumoniae* ATCC | 0.170752 |
| *Kluyveromyces marxianus* compared to *Moraxella catarrhalis* ATCC | <0.000001 |
| *Kluyveromyces marxianus* compared to *Escherichia coli* ATCC | <0.000001 |
| *Candida albicans* ATCC compared to *Candida glabrata* | <0.000001 |
| *Candida albicans* ATCC compared to *Candida krusei* | <0.000001 |
| *Candida albicans* ATCC compared to *Staphylococcus epidermidis* ATCC | 0.808098 |
| *Candida albicans* ATCC compared to *Enterococcus faecalis* ATCC | <0.000001 |
| *Candida albicans* ATCC compared to MSSA ATCC | 0.399741 |
| *Candida albicans* ATCC compared to MRSA ATCC | <0.000001 |
| *Candida albicans* ATCC compared to *Bacillus spizizenii* ATCC | <0.000001 |
| *Candida albicans* ATCC compared to *Proteus mirabilis* ATCC | <0.000001 |
| *Candida albicans* ATCC compared to *Enterobacter aerogenes* ATCC | <0.000001 |
| *Candida albicans* ATCC compared to *Salmonella enteritidis* ATCC | <0.000001 |
| *Candida albicans* ATCC compared to *Pseudomonas aeruginosa* ATCC | <0.000001 |
| *Candida albicans* ATCC compared to *Citrobacter freundii* ATCC | <0.000001 |
| *Candida albicans* ATCC compared to *Klebsiella pneumoniae* ATCC | 0.223873 |
| *Candida albicans* ATCC compared to *Moraxella catarrhalis* ATCC | <0.000001 |
| *Candida albicans* ATCC compared to *Escherichia coli* ATCC | <0.000001 |
| *Candida glabrata* compared to *Candida krusei* | <0.000001 |
| *Candida glabrata* compared to *Staphylococcus epidermidis* ATCC | 0.000956 |
| *Candida glabrata* compared to *Enterococcus faecalis* ATCC | <0.000001 |
| *Candida glabrata* compared to MSSA ATCC | 0.059364 |
| *Candida glabrata* compared to MRSA ATCC | 0.000168 |
| *Candida glabrata* compared to *Bacillus spizizenii* ATCC | <0.000001 |
| *Candida glabrata* compared to *Proteus mirabilis* ATCC | <0.000001 |
| *Candida glabrata* compared to *Enterobacter aerogenes* ATCC | <0.000001 |
| *Candida glabrata* compared to *Salmonella enteritidis* ATCC | <0.000001 |
| *Candida glabrata* compared to *Pseudomonas aeruginosa* ATCC | <0.000001 |
| *Candida glabrata* compared to *Citrobacter freundii* ATCC | <0.000001 |
| *Candida glabrata* compared to *Klebsiella pneumoniae* ATCC | 0.000031 |
| *Candida glabrata* compared to *Moraxella catarrhalis* ATCC | <0.000001 |
| *Candida glabrata* compared to *Escherichia coli* ATCC | <0.000001 |
| *Candida krusei* compared to *Staphylococcus epidermidis* ATCC | <0.000001 |
| *Candida krusei* compared to *Enterococcus faecalis* ATCC | <0.000001 |
| *Candida krusei* compared to MSSA ATCC | <0.000001 |
| *Candida krusei* compared to MRSA ATCC | 0.169116 |
| *Candida krusei* compared to *Bacillus spizizenii* ATCC | <0.000001 |
| *Candida krusei* compared to *Proteus mirabilis* ATCC | <0.000001 |
| *Candida krusei* compared to *Enterobacter aerogenes* ATCC | <0.000001 |
| *Candida krusei* compared to *Salmonella enteritidis* ATCC | <0.000001 |
| *Candida krusei* compared to *Pseudomonas aeruginosa* ATCC | 0.000310 |
| *Candida krusei* compared to *Citrobacter freundii* ATCC | <0.000001 |
| *Candida krusei* compared to *Klebsiella pneumoniae* ATCC | <0.000001 |
| *Candida krusei* compared to *Moraxella catarrhalis* ATCC | 0.000062 |
| *Candida krusei* compared to *Escherichia coli* ATCC | <0.000001 |
| *Staphylococcus epidermidis* ATCC compared to *Enterococcus faecalis* ATCC | <0.000001 |
| *Staphylococcus epidermidis* ATCC compared to MSSA ATCC | 0.594318 |
| *Staphylococcus epidermidis* ATCC compared to MRSA ATCC | <0.000001 |
| *Staphylococcus epidermidis* ATCC compared to *Bacillus spizizenii* ATCC | <0.000001 |
| *Staphylococcus epidermidis* ATCC compared to *Proteus mirabilis* ATCC | <0.000001 |
| *Staphylococcus epidermidis* ATCC compared to *Enterobacter aerogenes* ATCC | <0.000001 |
| *Staphylococcus epidermidis* ATCC compared to *Salmonella enteritidis* ATCC | <0.000001 |
| *Staphylococcus epidermidis* ATCC compared to *Pseudomonas aeruginosa* ATCC | <0.000001 |
| *Staphylococcus epidermidis* ATCC compared to *Citrobacter freundii* ATCC | <0.000001 |
| *Staphylococcus epidermidis* ATCC compared to *Klebsiella pneumoniae* ATCC | 0.264942 |
| *Staphylococcus epidermidis* ATCC compared to *Moraxella catarrhalis* ATCC | <0.000001 |
| *Staphylococcus epidermidis* ATCC compared to *Escherichia coli* ATCC | <0.000001 |
| *Enterococcus faecalis* ATCC compared to MSSA ATCC | <0.000001 |
| *Enterococcus faecalis* ATCC compared to MRSA ATCC | 0.029654 |
| *Enterococcus faecalis* ATCC compared to *Bacillus spizizenii* ATCC | <0.000001 |
| *Enterococcus faecalis* ATCC compared to *Proteus mirabilis* ATCC | 0.273961 |
| *Enterococcus faecalis* ATCC compared to *Enterobacter aerogenes* ATCC | 0.005731 |
| *Enterococcus faecalis* ATCC compared to *Salmonella enteritidis* ATCC | 0.000023 |
| *Enterococcus faecalis* ATCC compared to *Pseudomonas aeruginosa* ATCC | 0.753577 |
| *Enterococcus faecalis* ATCC compared to *Citrobacter freundii* ATCC | 0.024527 |
| *Enterococcus faecalis* ATCC compared to *Klebsiella pneumoniae* ATCC | <0.000001 |
| *Enterococcus faecalis* ATCC compared to *Moraxella catarrhalis* ATCC | 0.440614 |
| *Enterococcus faecalis* ATCC compared to *Escherichia coli* ATCC | 0.686266 |
| MSSA ATCC compared to MRSA ATCC | 0.000037 |
| MSSA ATCC compared to *Bacillus spizizenii* ATCC | <0.000001 |
| MSSA ATCC compared to *Proteus mirabilis* ATCC | <0.000001 |
| MSSA ATCC compared to *Enterobacter aerogenes* ATCC | <0.000001 |
| MSSA ATCC compared to *Salmonella enteritidis* ATCC | <0.000001 |
| MSSA ATCC compared to *Pseudomonas aeruginosa* ATCC | <0.000001 |
| MSSA ATCC compared to *Citrobacter freundii* ATCC | <0.000001 |
| MSSA ATCC compared to *Klebsiella pneumoniae* ATCC | 0.141919 |
| MSSA ATCC compared to *Moraxella catarrhalis* ATCC | <0.000001 |
| MSSA ATCC compared to *Escherichia coli* ATCC | <0.000001 |
| MRSA ATCC compared to *Bacillus spizizenii* ATCC | <0.000001 |
| MRSA ATCC compared to *Proteus mirabilis* ATCC | 0.001639 |
| MRSA ATCC compared to *Enterobacter aerogenes* ATCC | 0.000002 |
| MRSA ATCC compared to *Salmonella enteritidis* ATCC | <0.000001 |
| MRSA ATCC compared to *Pseudomonas aeruginosa* ATCC | 0.085947 |
| MRSA ATCC compared to *Citrobacter freundii* ATCC | 0.000017 |
| MRSA ATCC compared to *Klebsiella pneumoniae* ATCC | <0.000001 |
| MRSA ATCC compared to *Moraxella catarrhalis* ATCC | 0.122789 |
| MRSA ATCC compared to *Escherichia coli* ATCC | 0.010245 |
| *Bacillus spizizenii* ATCC compared to *Proteus mirabilis* ATCC | <0.000001 |
| *Bacillus spizizenii* ATCC compared to *Enterobacter aerogenes* ATCC | <0.000001 |
| *Bacillus spizizenii* ATCC compared to *Salmonella enteritidis* ATCC | <0.000001 |
| *Bacillus spizizenii* ATCC compared to *Pseudomonas aeruginosa* ATCC | <0.000001 |
| *Bacillus spizizenii* ATCC compared to *Citrobacter freundii* ATCC | <0.000001 |
| *Bacillus spizizenii* ATCC compared to *Klebsiella pneumoniae* ATCC | 0.000001 |
| *Bacillus spizizenii* ATCC compared to *Moraxella catarrhalis* ATCC | <0.000001 |
| *Bacillus spizizenii* ATCC compared to *Escherichia coli* ATCC | <0.000001 |
| *Proteus mirabilis* ATCC compared to *Enterobacter aerogenes* ATCC | 0.107392 |
| *Proteus mirabilis* ATCC compared to *Salmonella enteritidis* ATCC | 0.002348 |
| *Proteus mirabilis* ATCC compared to *Pseudomonas aeruginosa* ATCC | 0.188112 |
| *Proteus mirabilis* ATCC compared to *Citrobacter freundii* ATCC | 0.280240 |
| *Proteus mirabilis* ATCC compared to *Klebsiella pneumoniae* ATCC | <0.000001 |
| *Proteus mirabilis* ATCC compared to *Moraxella catarrhalis* ATCC | 0.059122 |
| *Proteus mirabilis* ATCC compared to *Escherichia coli* ATCC | 0.478179 |
| *Enterobacter aerogenes* ATCC compared to *Salmonella enteritidis* ATCC | 0.145333 |
| *Enterobacter aerogenes* ATCC compared to *Pseudomonas aeruginosa* ATCC | 0.004296 |
| *Enterobacter aerogenes* ATCC compared to *Citrobacter freundii* ATCC | 0.563406 |
| *Enterobacter aerogenes* ATCC compared to *Klebsiella pneumoniae* ATCC | <0.000001 |
| *Enterobacter aerogenes* ATCC compared to *Moraxella catarrhalis* ATCC | 0.000273 |
| *Enterobacter aerogenes* ATCC compared to *Escherichia coli* ATCC | 0.017078 |
| *Salmonella enteritidis* ATCC compared to *Pseudomonas aeruginosa* ATCC | 0.000025 |
| *Salmonella enteritidis* ATCC compared to *Citrobacter freundii* ATCC | 0.038555 |
| *Salmonella enteritidis* ATCC compared to *Klebsiella pneumoniae* ATCC | <0.000001 |
| *Salmonella enteritidis* ATCC compared to *Moraxella catarrhalis* ATCC | <0.000001 |
| *Salmonella enteritidis* ATCC compared to *Escherichia coli* ATCC | 0.000111 |
| *Pseudomonas aeruginosa* ATCC compared to *Citrobacter freundii* ATCC | 0.017290 |
| *Pseudomonas aeruginosa* ATCC compared to *Klebsiella pneumoniae* ATCC | <0.000001 |
| *Pseudomonas aeruginosa* ATCC compared to *Moraxella catarrhalis* ATCC | 0.705969 |
| *Pseudomonas aeruginosa* ATCC compared to *Escherichia coli* ATCC | 0.492334 |
| *Citrobacter freundii* ATCC compared to *Klebsiella pneumoniae* ATCC | <0.000001 |
| *Citrobacter freundii* ATCC compared to *Moraxella catarrhalis* ATCC | 0.001741 |
| *Citrobacter freundii* ATCC compared to *Escherichia coli* ATCC | 0.063245 |
| *Klebsiella pneumoniae* ATCC compared to *Moraxella catarrhalis* ATCC | <0.000001 |
| *Klebsiella pneumoniae* ATCC compared to *Escherichia coli* ATCC | <0.000001 |
| *Moraxella catarrhalis* ATCC compared to *Escherichia coli* ATCC | 0.230218 |

## Rusult summary

Comparing the results from the previous tables (Table 5), it can be noted that silicon dioxide improved the chances of distinguishing one bacteria from another at 285-1100 nm. That increased the overall results. It also decreased the odds of distinguishing one yeast from another. At 285-700 nm silicon dioxide did not have a significant impact on the results, but it can be noted that the chances of distinguishing one bacteria of another increased at this wavelength interval in comparison with 285-1100 nm.

**Table 5.** Summary of all comparisons.

| Compared types of cultures | Comparisons without silicon dioxide at 285-1100 nm | Comparisons with silicon dioxide at 285-1100 nm | Comparisons without silicon dioxide at 285-700 nm | Comparisons with silicon dioxide at 285-700 nm |
| --- | --- | --- | --- | --- |
| Overall | 99/136 (72.8%) | 109/136 (81.1%) | 106/136 (77.9%) | 107/136 (78.7%) |
| Yeast compared to bacteria | 52/52 (100%) | 52/52 (100%) | 44/52 (84.6%) | 44/52 (84.6%) |
| Bacteria compared to bacteria | 41/78 (52.6%) | 54/78 (69.2%) | 56/78 (71.8%) | 58/78 (74.6%) |
| Yeast compared to yeast | 5/6 (83.3%) | 3/6 (50%) | 6/6 (100%) | 5/6 (83.3%) |

### Silicon dioxide effect on average absorbance

Silicon dioxide decreased the average light absorption of most yeast in a measurement interval of 285-1100 nm, assuming a significant change of 0.03 absorbance units, and increased the average light absorption of most bacteria in the same measurement interval (Table 6). For *Candida glabrata* the average light absorption did not have a significant decrease. For *Staphylococcus epidermidis* the average absorbance adding silicon dioxide decreased, but for *Enterococcus faecalis*, MSSA, *Proteus mirabilis* and *Escherichia coli* it had no significant changes.

**Table 6.** Effects of silicon dioxide on the average absorbance of different microorganisms.

| Microorganisms | Average absorbance without silicon dioxide | Average absorbance with silicon dioxide | Silicon dioxide effect on average absorbance |
| --- | --- | --- | --- |
| *Kluyveromyces marxianus* | 1.262251 | 1.158944 | Decrease |
| *Candida albicans* ATCC | 1.227039 | 1.135811 | Decrease |
| *Candida glabrata* | 1.180000 | 1.168283 | Decrease |
| *Candida krusei* | 1.194647 | 1.134603 | Decrease |
| *Staphylococcus epidermidis* ATCC | 0.946494 | 0.895001 | Decrease |
| *Enterococcus faecalis* ATCC | 0.949259 | 0.942527 | Did not alter |
| MSSA ATCC | 0.793623 | 0.812563 | Increase |
| MRSA ATCC | 0.830097 | 0.886495 | Increase |
| *Bacillus spizizenii* ATCC | 0.715224 | 0.745946 | Increase |
| *Proteus mirabilis* ATCC | 0.935026 | 0.957902 | Increase |
| *Enterobacter aerogenes* ATCC | 0.938885 | 0.982013 | Increase |
| *Salmonella enteritidis* ATCC | 0.941685 | 1.006821 | Increase |
| *Pseudomonas aeruginosa* ATCC | 0.892804 | 0.930857 | Increase |
| *Citrobacter freundii* ATCC | 0.930903 | 0.991864 | Increase |
| *Klebsiella pneumoniae* ATCC | 0.661763 | 0.837743 | Increase |
| *Moraxella catarrhalis* ATCC | 0.897066 | 0.935586 | Increase |
| *Escherichia coli* ATCC | 0.900817 | 0.905401 | Did not alter |

### Peculiarities of microorganism graphs

Analyzing the average graphs of yeast, Gram positive and Gram negative bacteria (Fig 1), it is possible to find certain characteristics that correspond to the type of microorganism that is being analyzed by the spectrophotometer. Graphs are shown from highest light absorption going to the lowest. Such order is maintained because a spectrophotometer presents its measured absorbance units starting at a higher wavelength and going to a lower. All yeast presented a higher light absorption than Gram positive and Gram negative bacteria at near infrared wavelengths. Light absorption for bacteria increased at a higher rate than that of yeast. And at some point the light absorbance units of bacteria surpassed the light absorbance units of yeast. At about 305 nm all microorganisms had a drastic spike in the increase of light absorption. It was more noticeable for bacteria than it was for yeast. These criteria could be used to distinguish bacteria from yeast. For Gram negative bacteria the spike at the end of the measurement was larger than it was for Gram positive bacteria. This criteria could be used to distinguish Gram negative bacteria from Gram positive bacteria.

**Fig 1.**
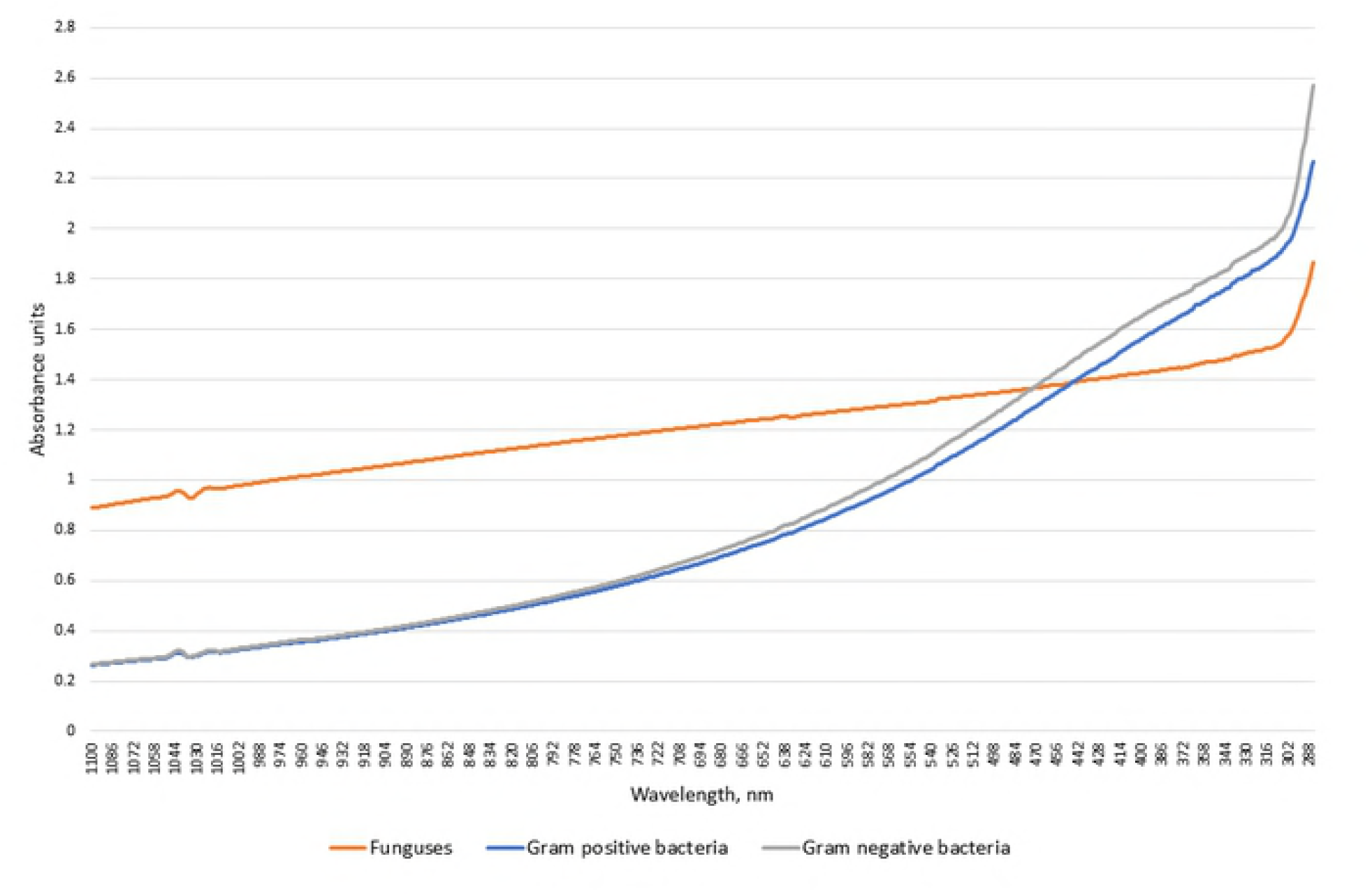
Average light absorption of yeast, Gram positive and Gram negative bacteria at 285-1100 nm without silicon dioxide.

Adding silicon dioxide, the characteristics, that were specific in distinguishing bacteria from yeast without silicon dioxide, remained the same (Fig 2). But silicon dioxide did produce alteration in microorganism graphs. Adding silicon dioxide, the average light absorption of all yeast deceased (Fig 3, Fig 4, Fig 5, Fig 7), except for *Candida glabrata*, for whom it stayed almost the same (Fig 6). For all bacteria absorbance increased at the near infrared wavelengths and for several bacteria it increased at visible light (380-700 nm) and for some even at ultraviolet (<380 nm) wavelengths (Fig 9–22, Table 7). The smallest increase of absorbance at near infrared wavelengths was for *Staphylococcus epidermidis* which was barely noticeable from 1100 nm going down till 900 nm (Fig 9). *Klebsiella pneumoniae* had the most noticeable increase of absorbance when silicon dioxide was added. The absorbance increased during the whole measurement (Fig 20). For *Salmonella enteritidis* the increase of absorbance, when silicon dioxide was added, was during almost the whole measurement (Fig 17).

**Fig 2.**
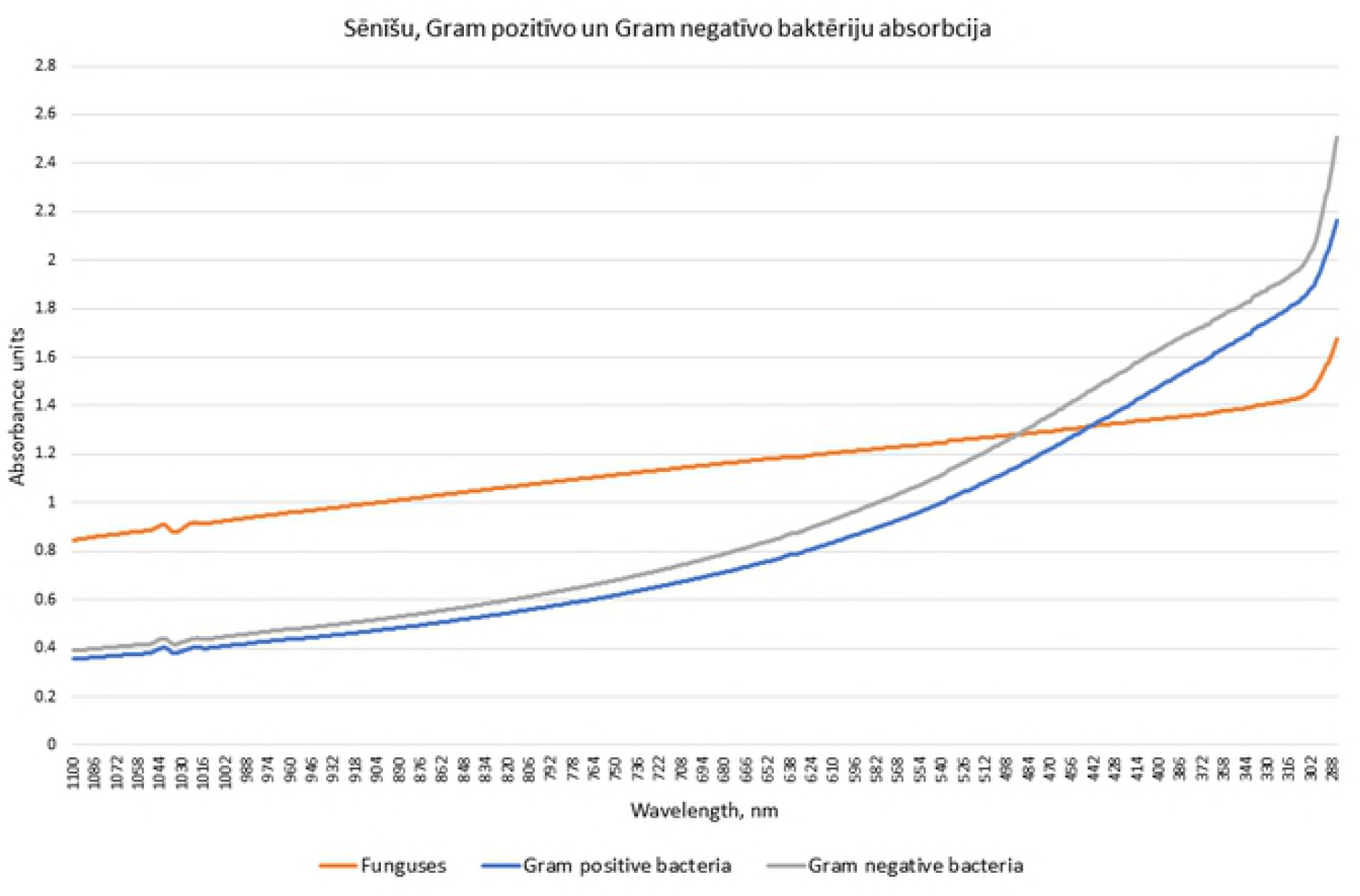
Average light absorption of all examined yeast, Gram positive and Gram negative bacteria at 285-1100 nm with silicon dioxide.

**Fig 3.**
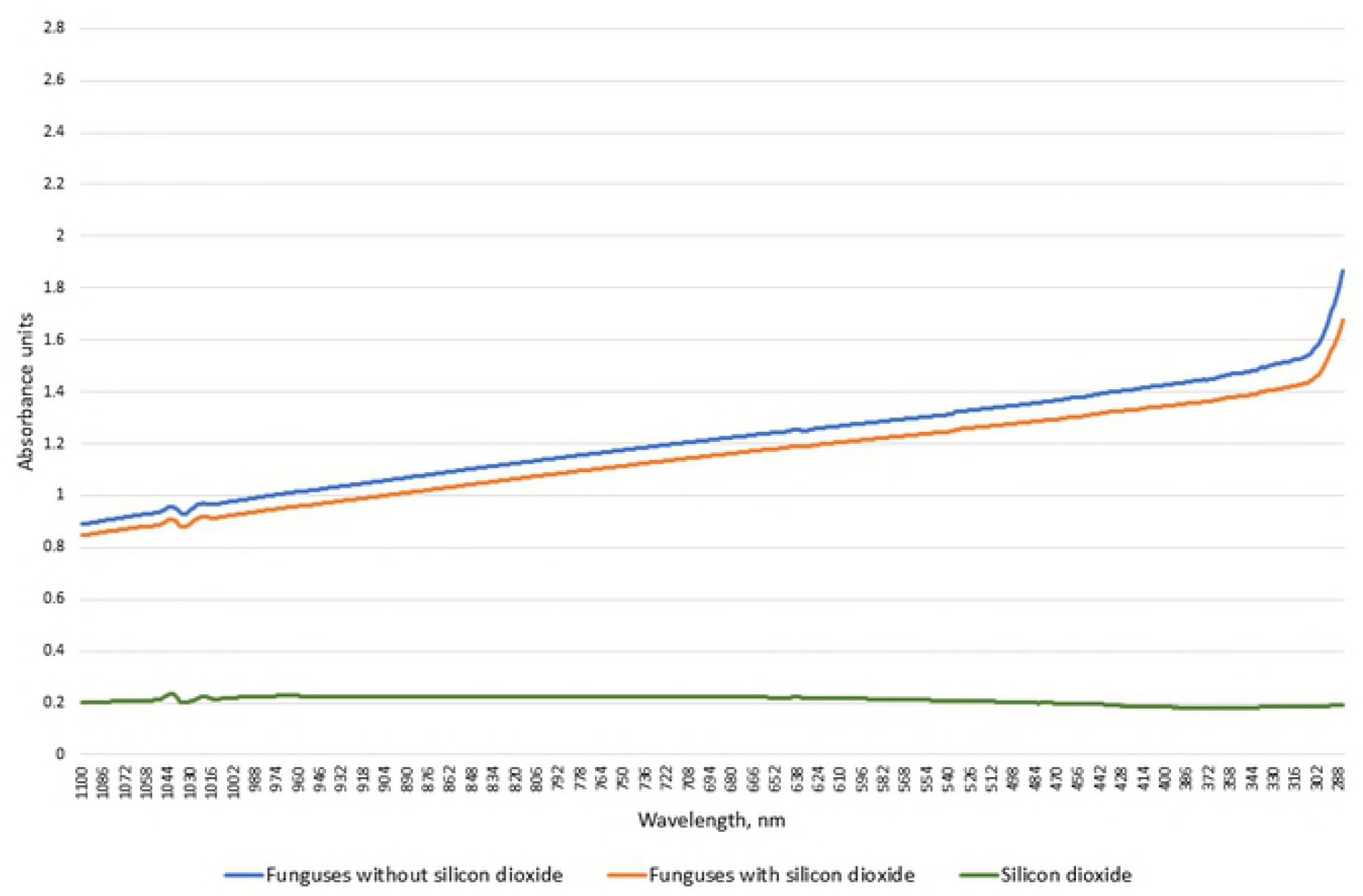
Average light absorption of all examined yeast without and with silicon dioxide.

**Fig 4.**
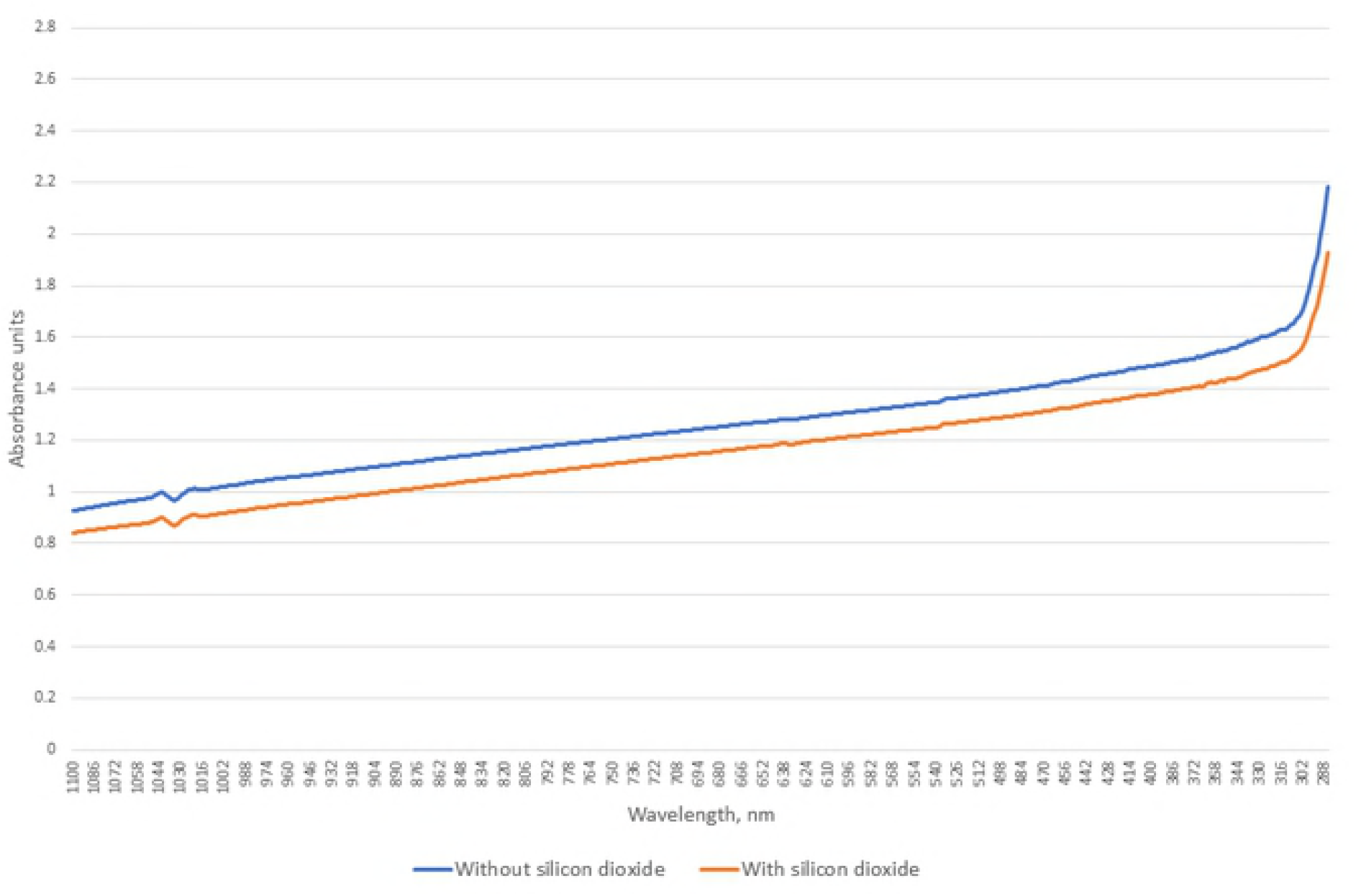
Light absorption of *Kluyveromyces marxianus* with and without silicon dioxide.

**Fig 5.**
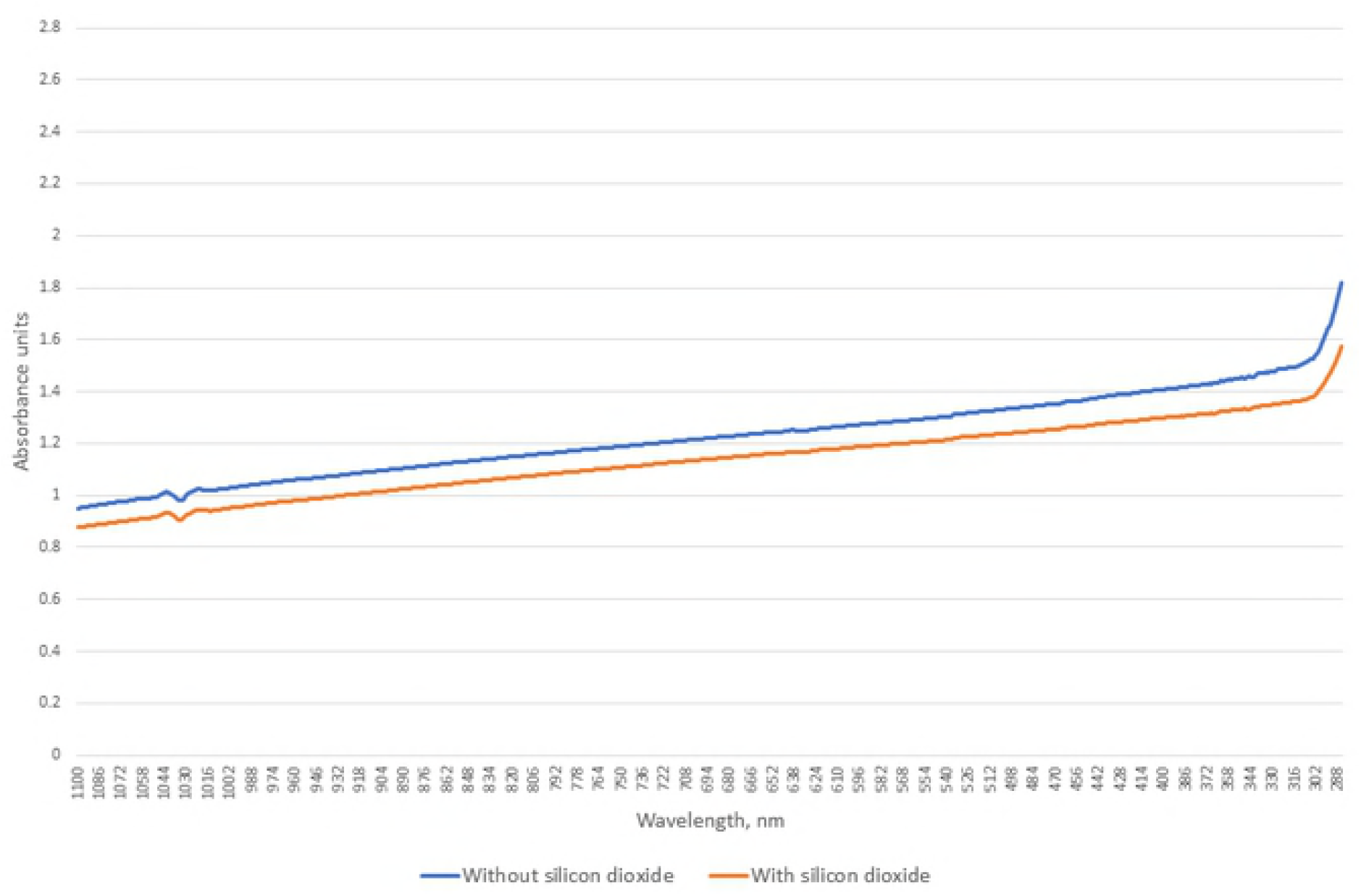
Light absorption of *Candida albicans* with and without silicon dioxide.

**Fig 6.**
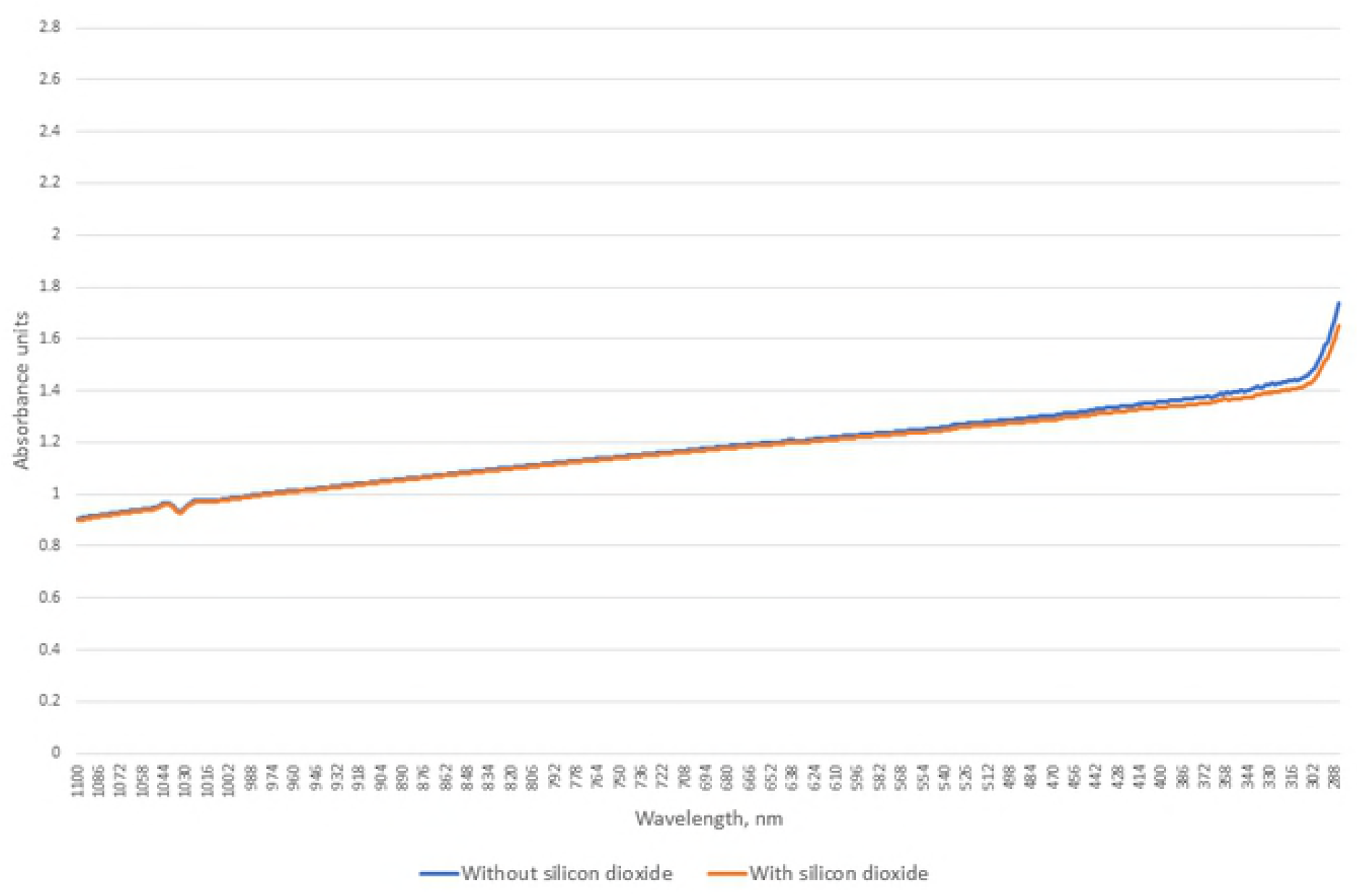
Light absorption of *Candida glabrata* with and without silicon dioxide.

**Fig 7.**
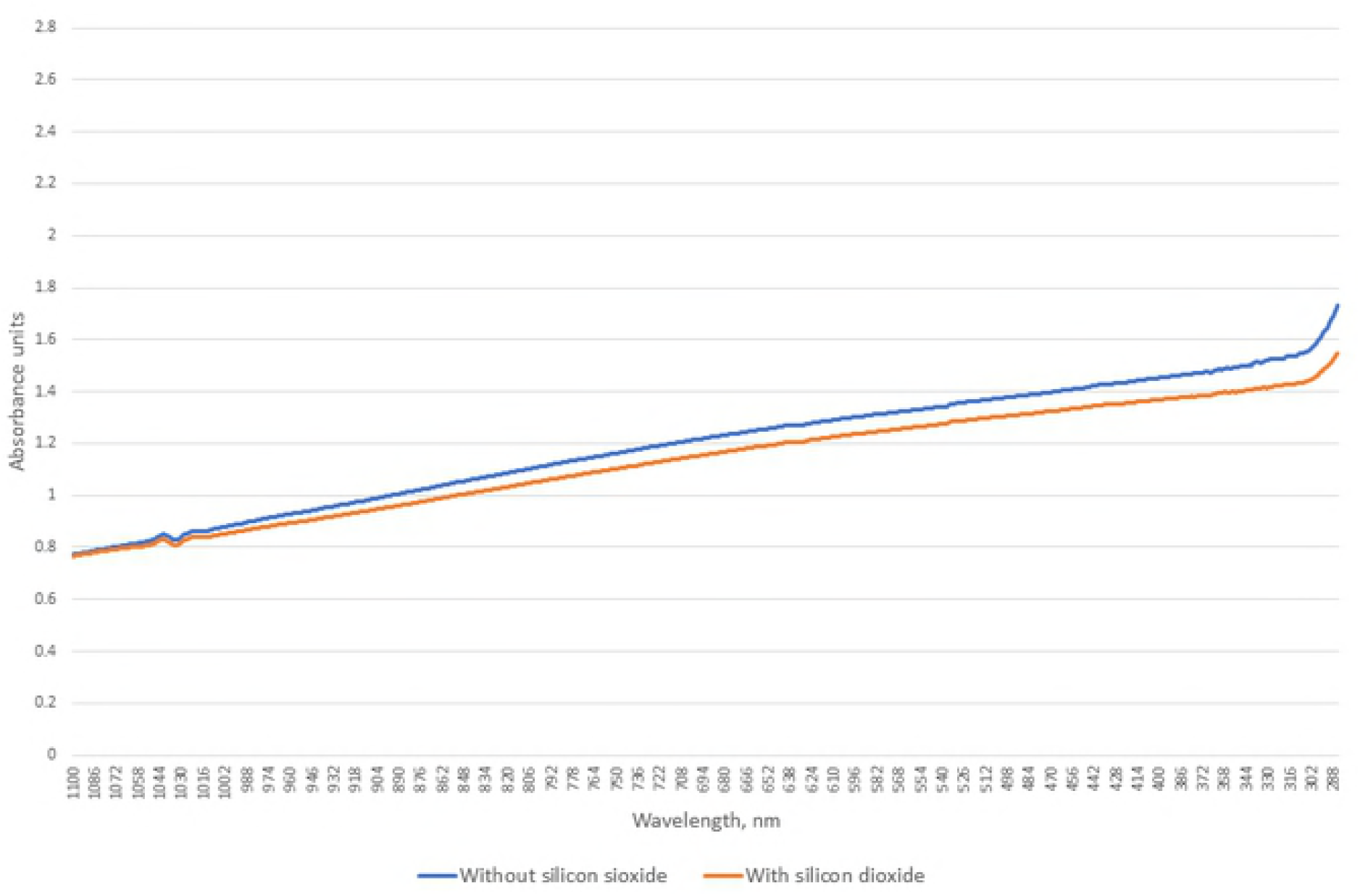
Light absorption of *Candida krusei* with and without silicon dioxide.

**Fig 8.**
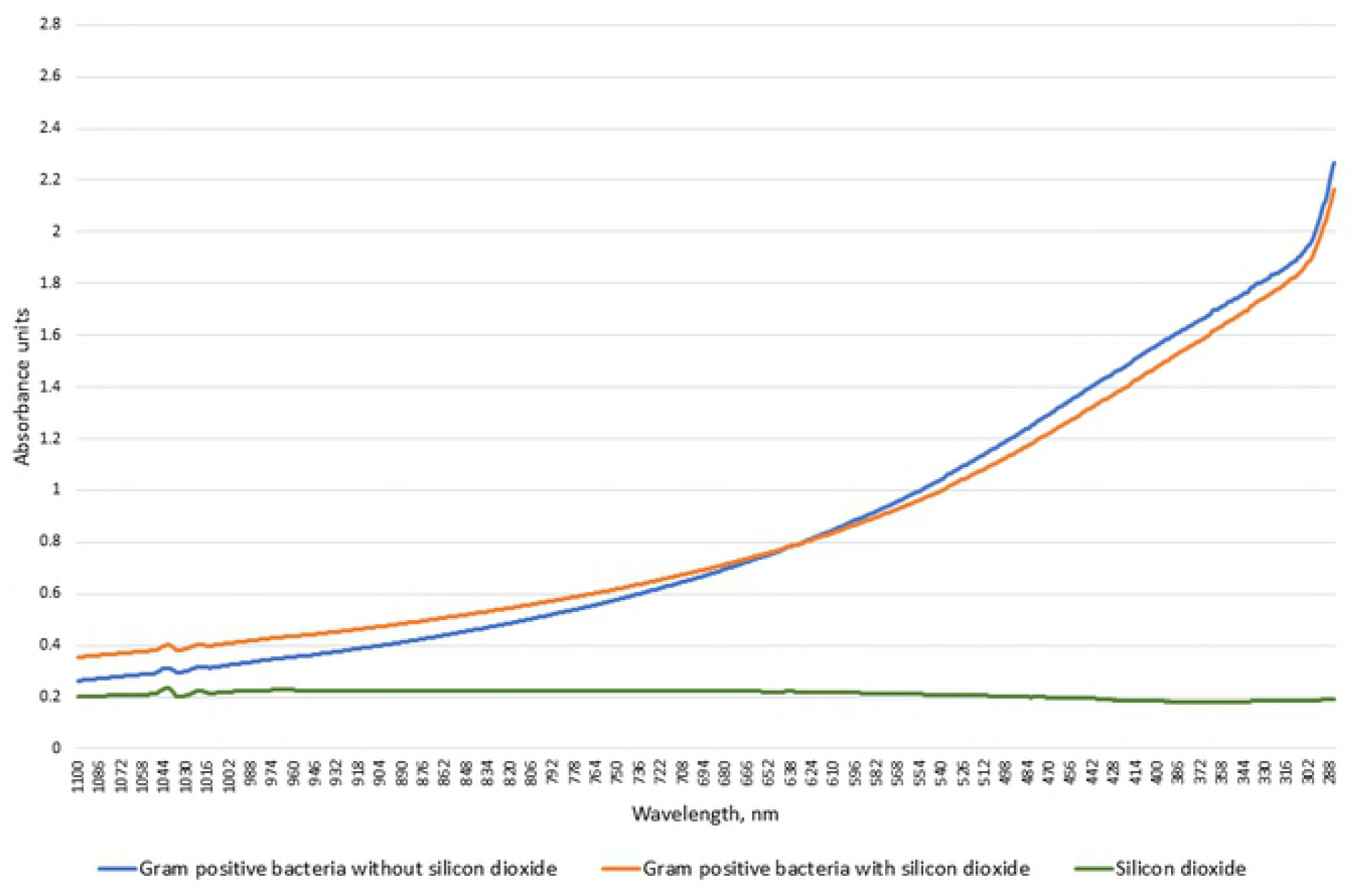
Average light absorption of all examined Gram positive bacteria without and with silicon dioxide.

**Fig 9.**
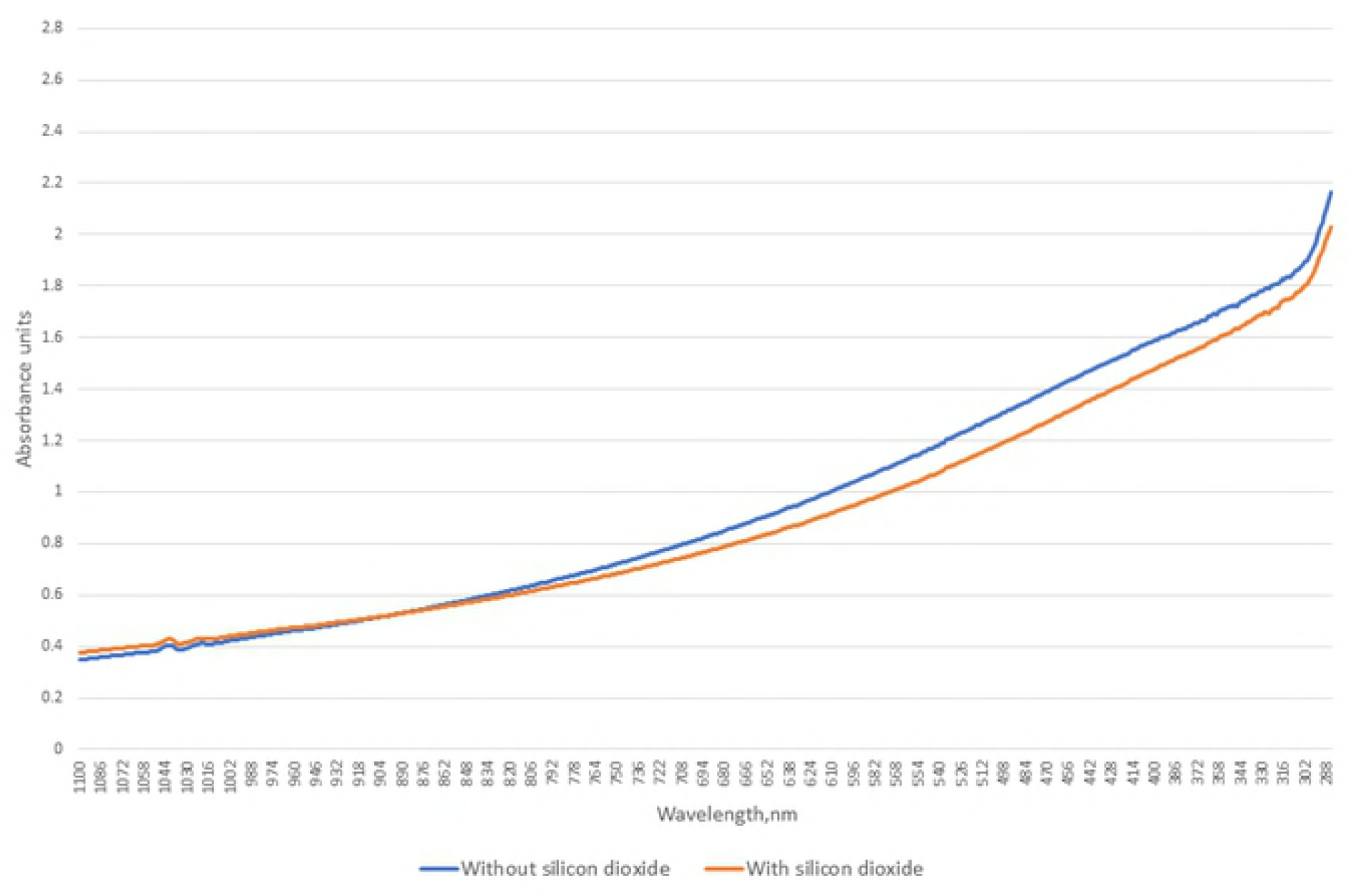
Light absorption of *Staphylococcus epidermidis* with and without silicon dioxide.

**Fig 10.**
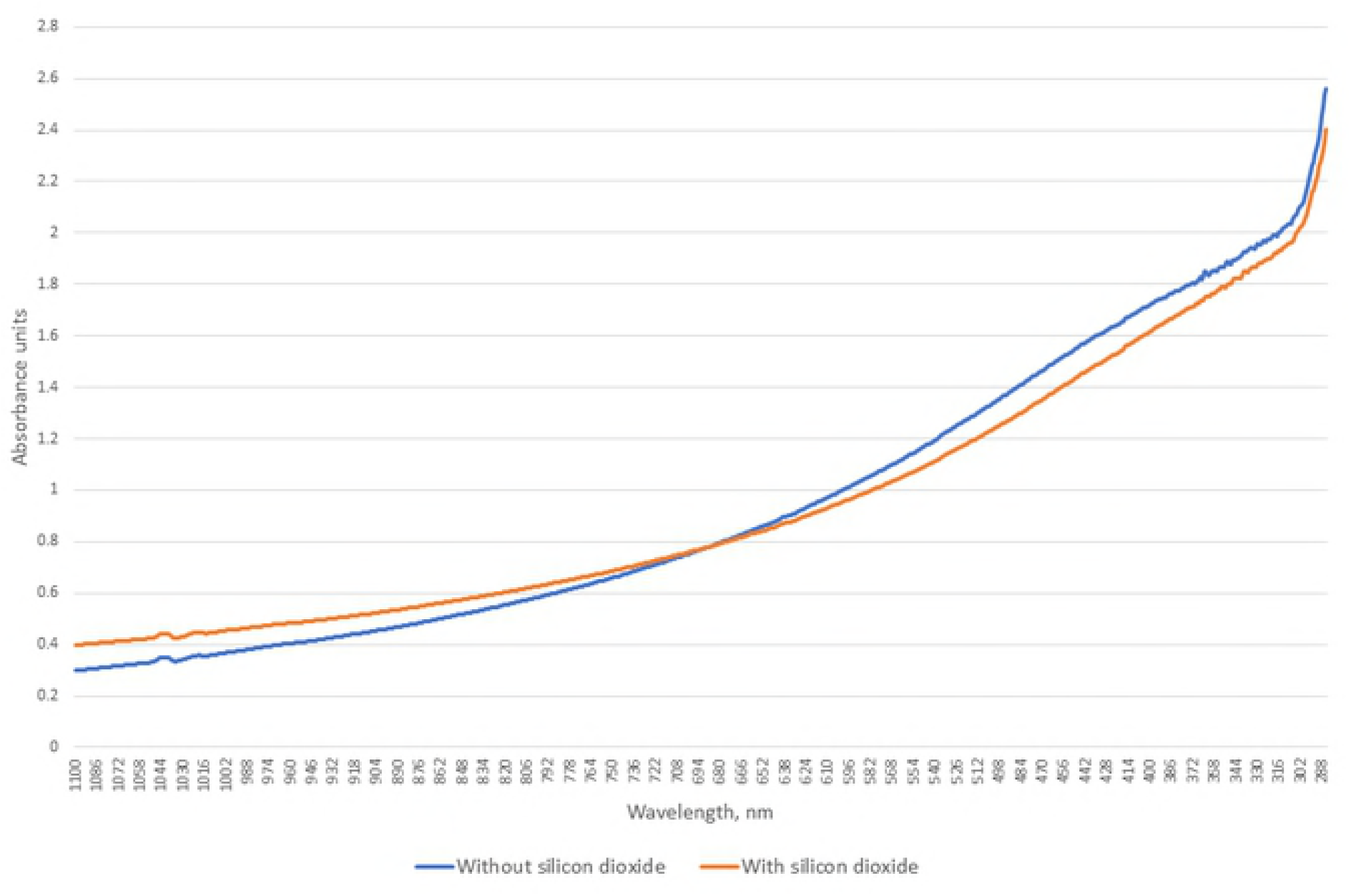
Light absorption of *Enterococcus faecalis* with and without silicon dioxide.

**Fig 11.**
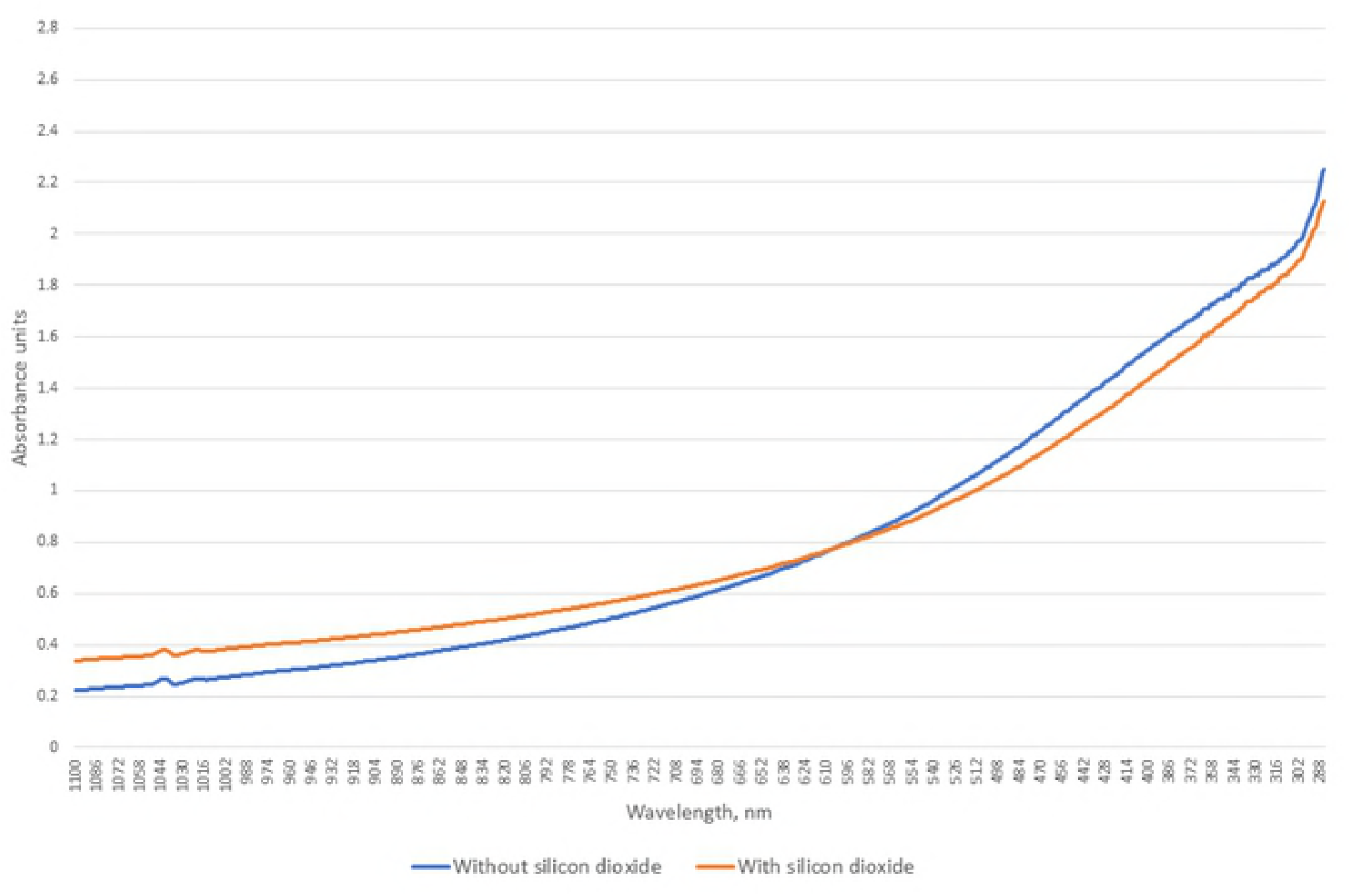
Light absorption of MSSA with and without silicon dioxide.

**Fig 12.**
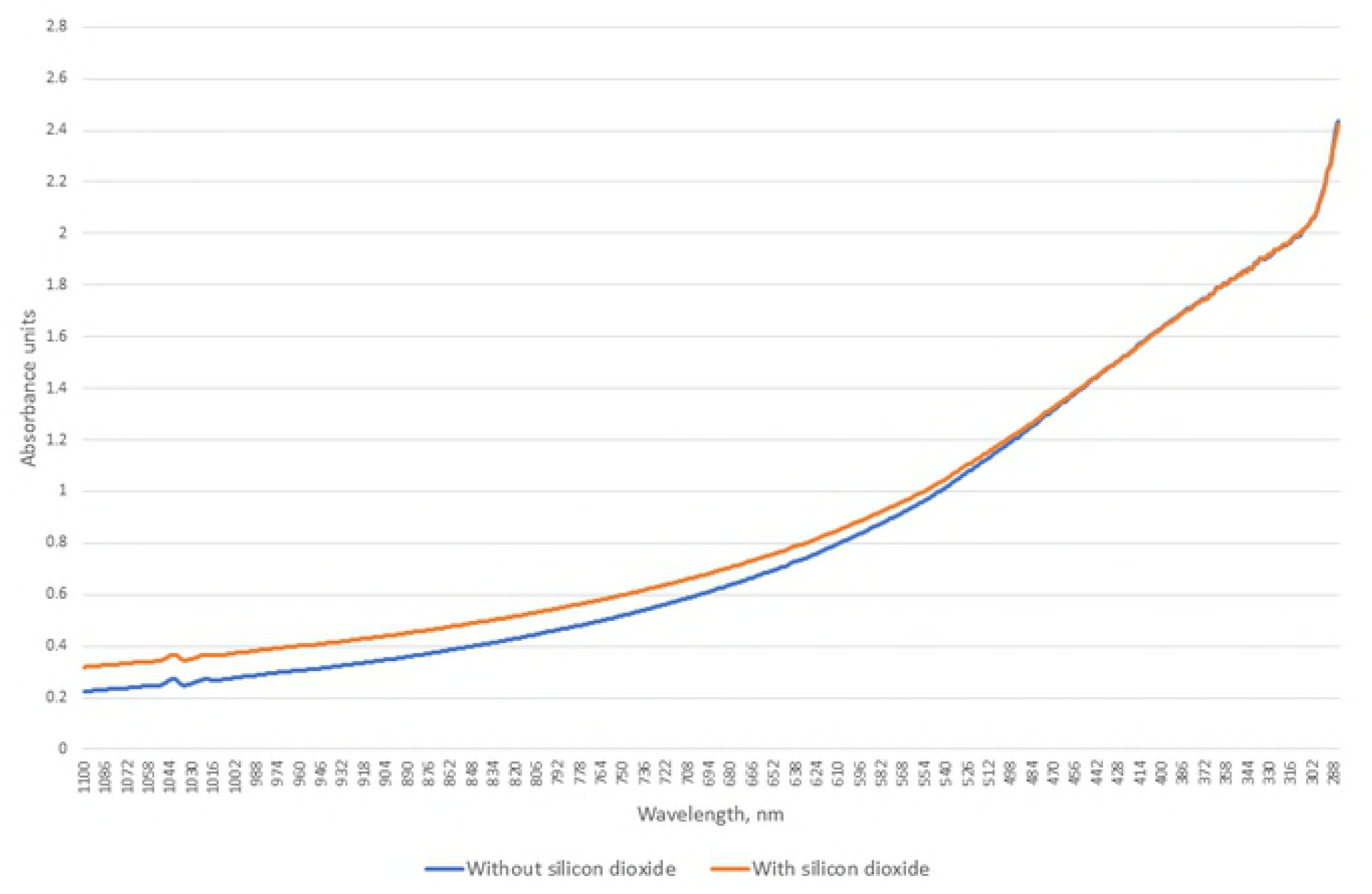
Light absorption of MRSA with and without silicon dioxide.

**Fig 13.**
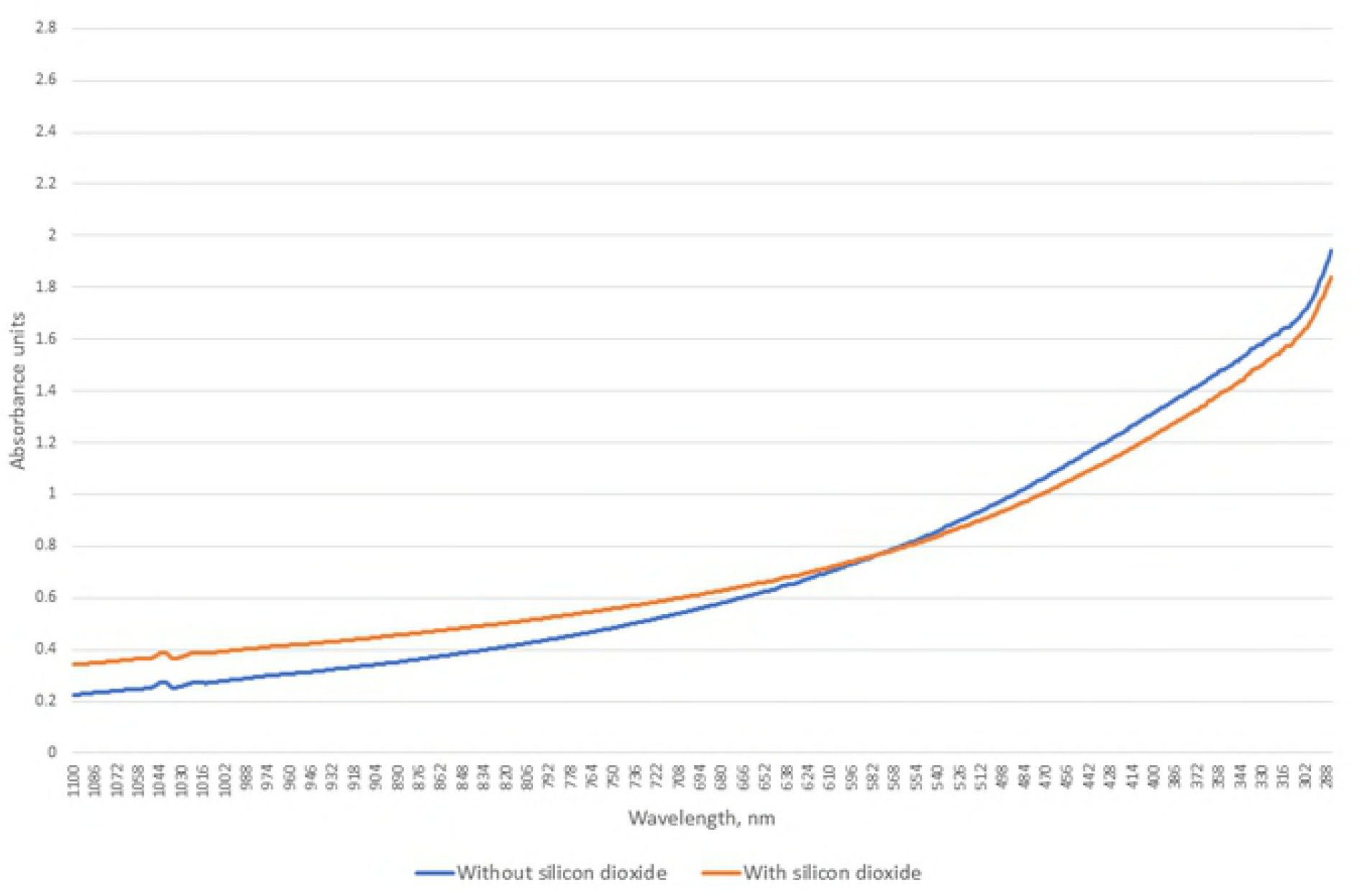
Light absorption of *Bacillus spizizenii* with and without silicon dioxide.

**Fig 14.**
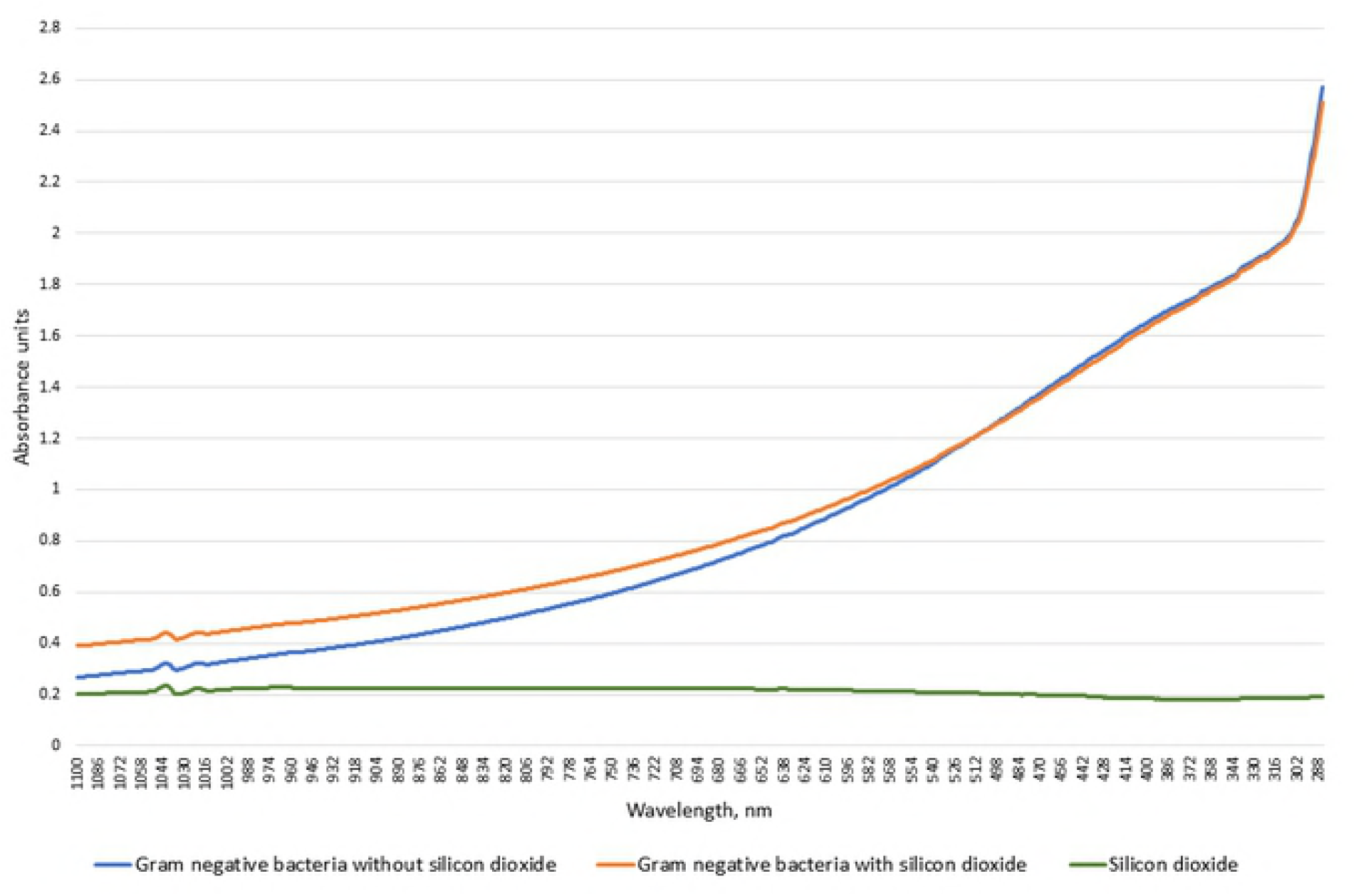
Average light absorption of all examined Gram negative bacteria without and with silicon dioxide.

**Fig 15.**
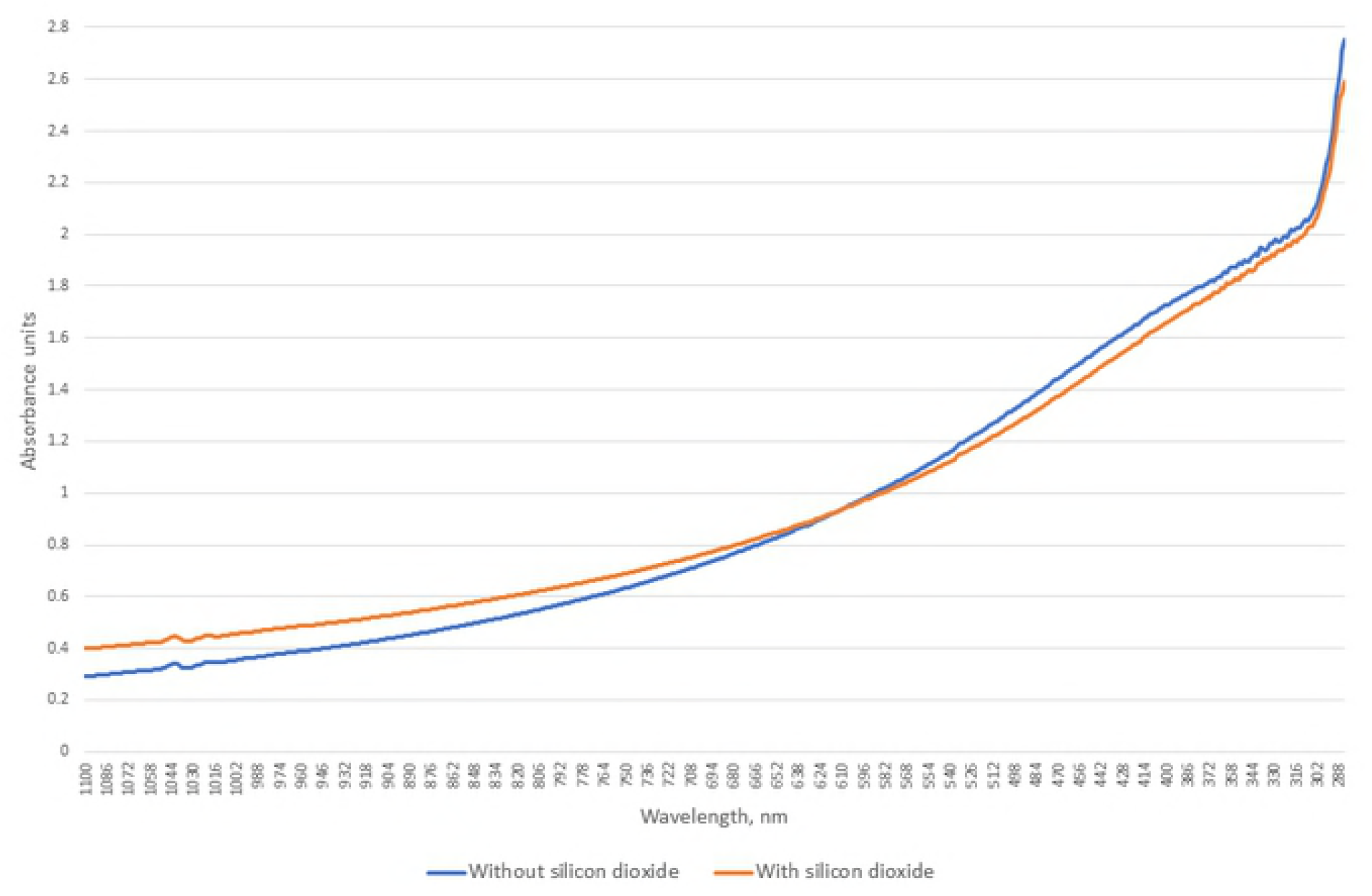
Light absorption of *Proteus mirabilis* with and without silicon dioxide.

**Fig 16.**
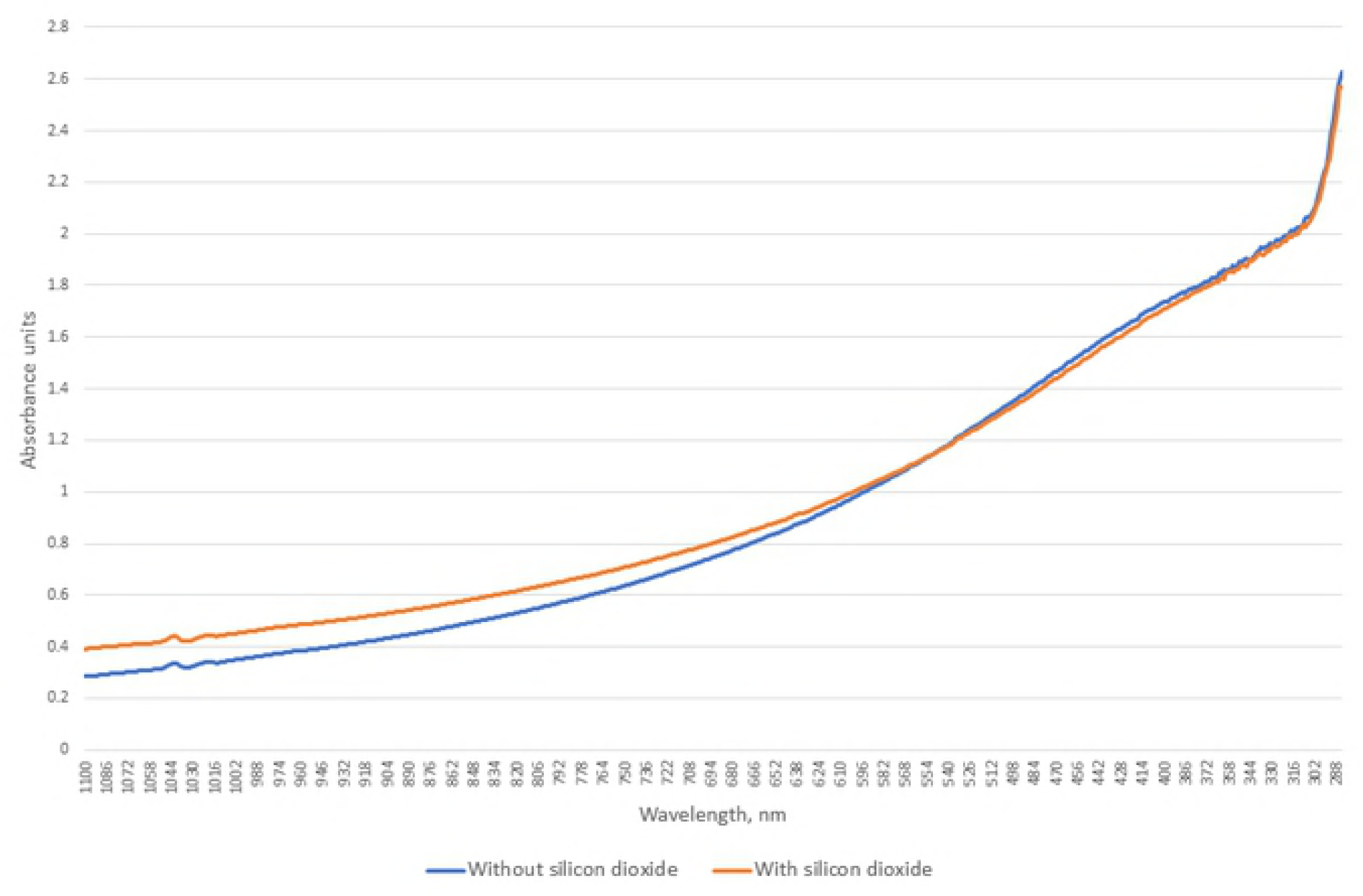
Light absorption of *Enterobacter aerogenes* with and without silicon dioxide.

**Fig 17.**
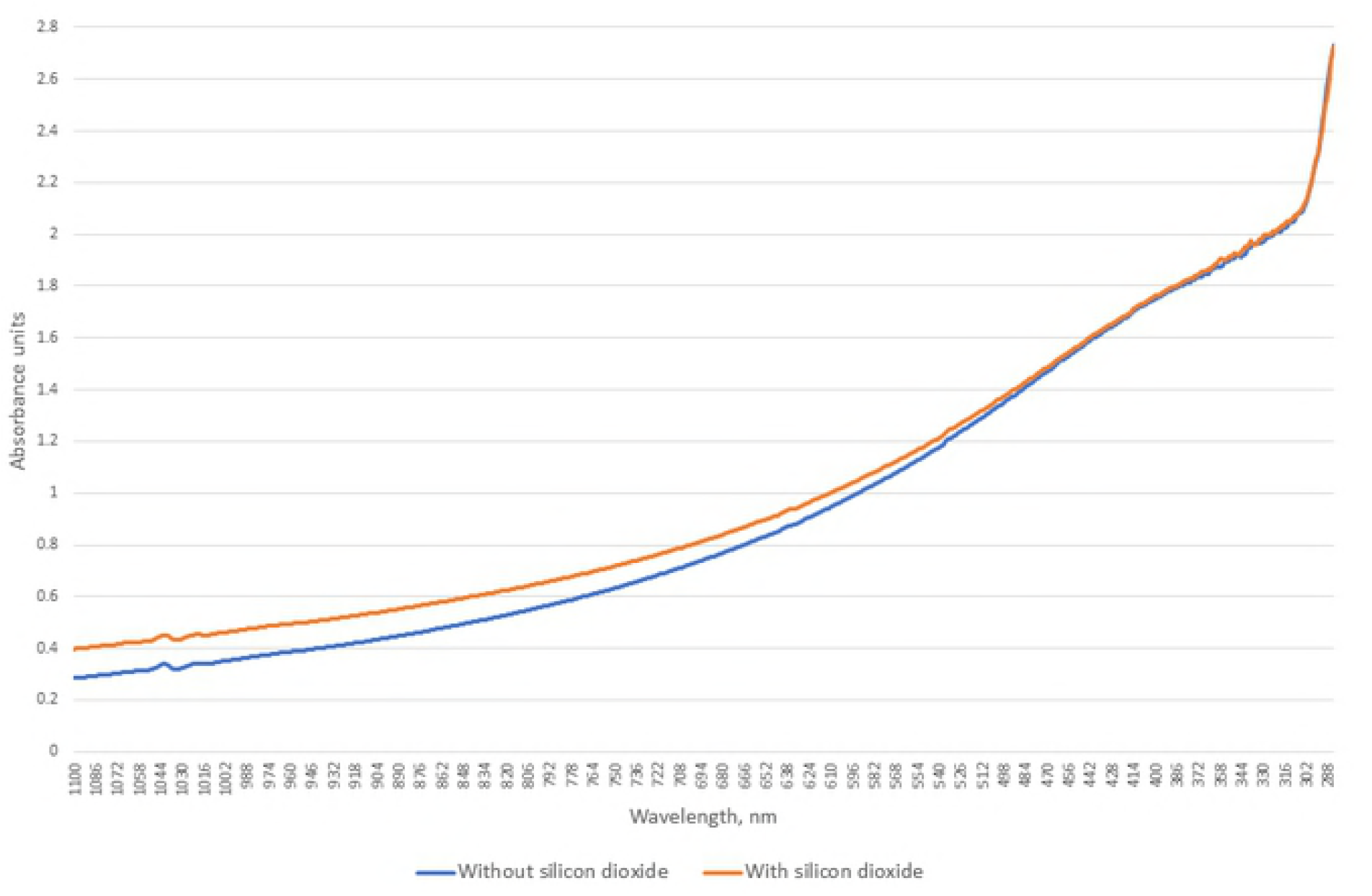
Light absorption of *Salmonella enteritidis* with and without silicon dioxide.

**Fig 18.**
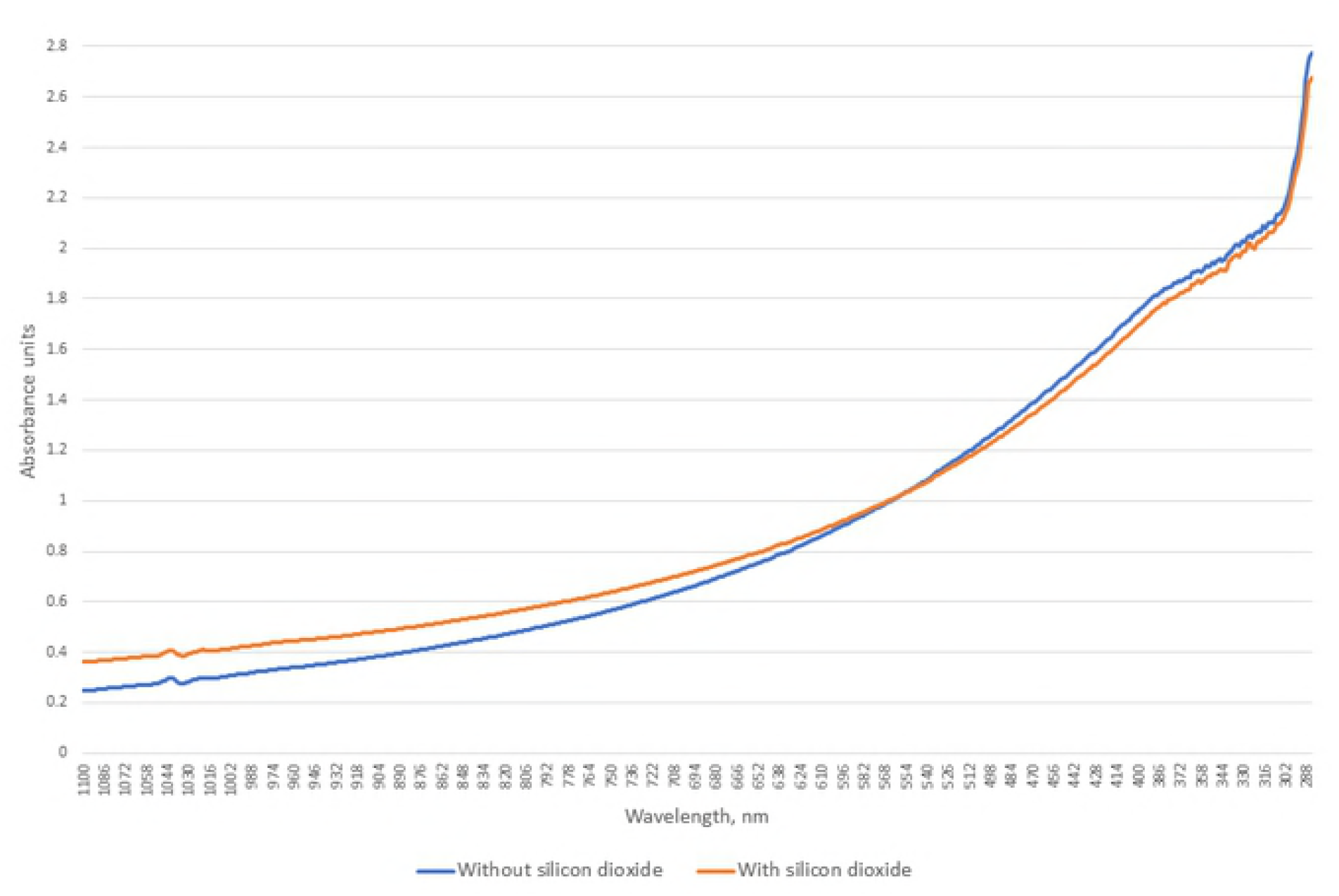
Light absorption of *Pseudomonas aeruginosa* with and without silicon dioxide.

**Fig 19.**
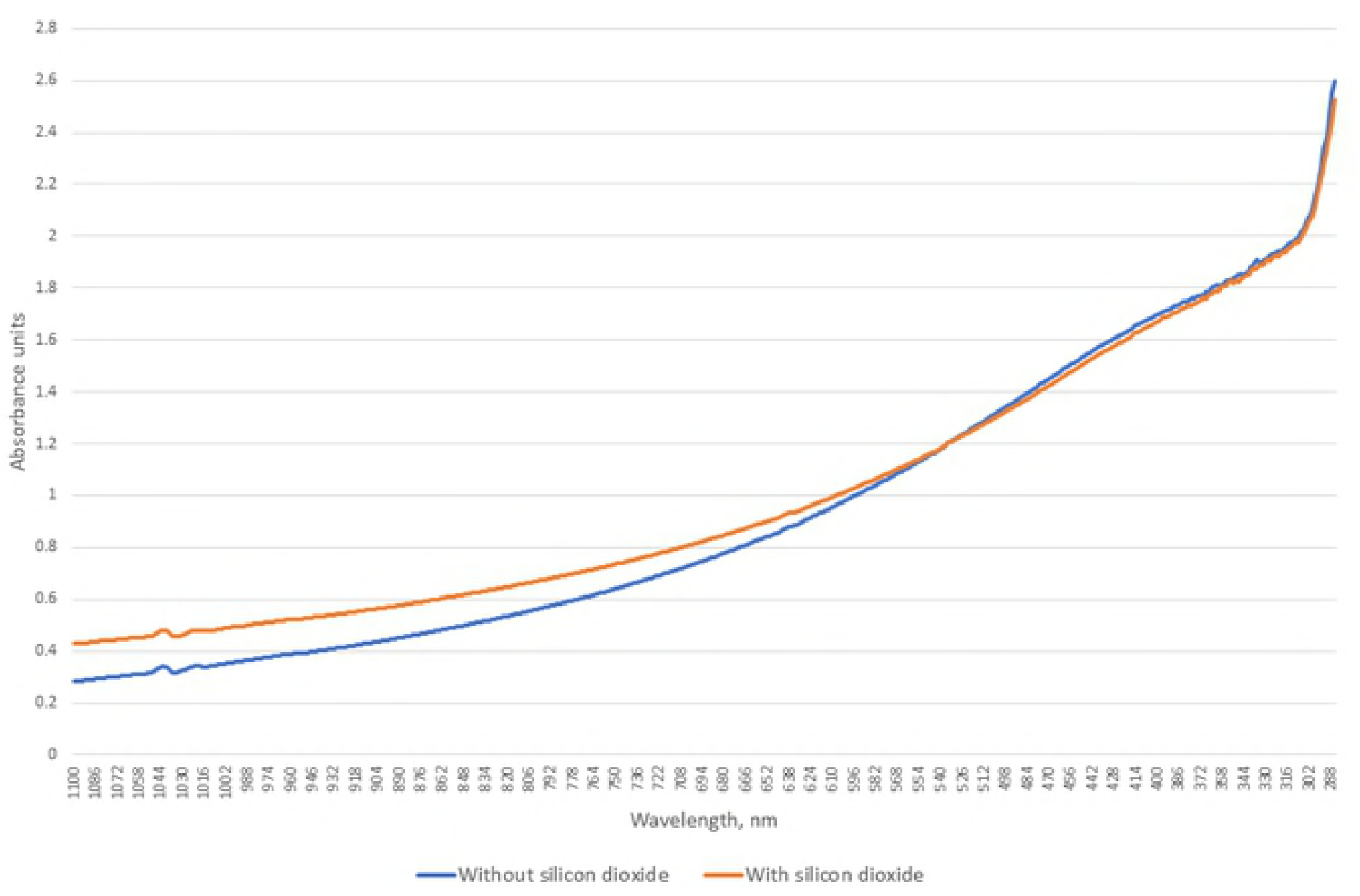
Light absorption of *Citrobacter freundii* with and without silicon dioxide.

**Fig 20.**
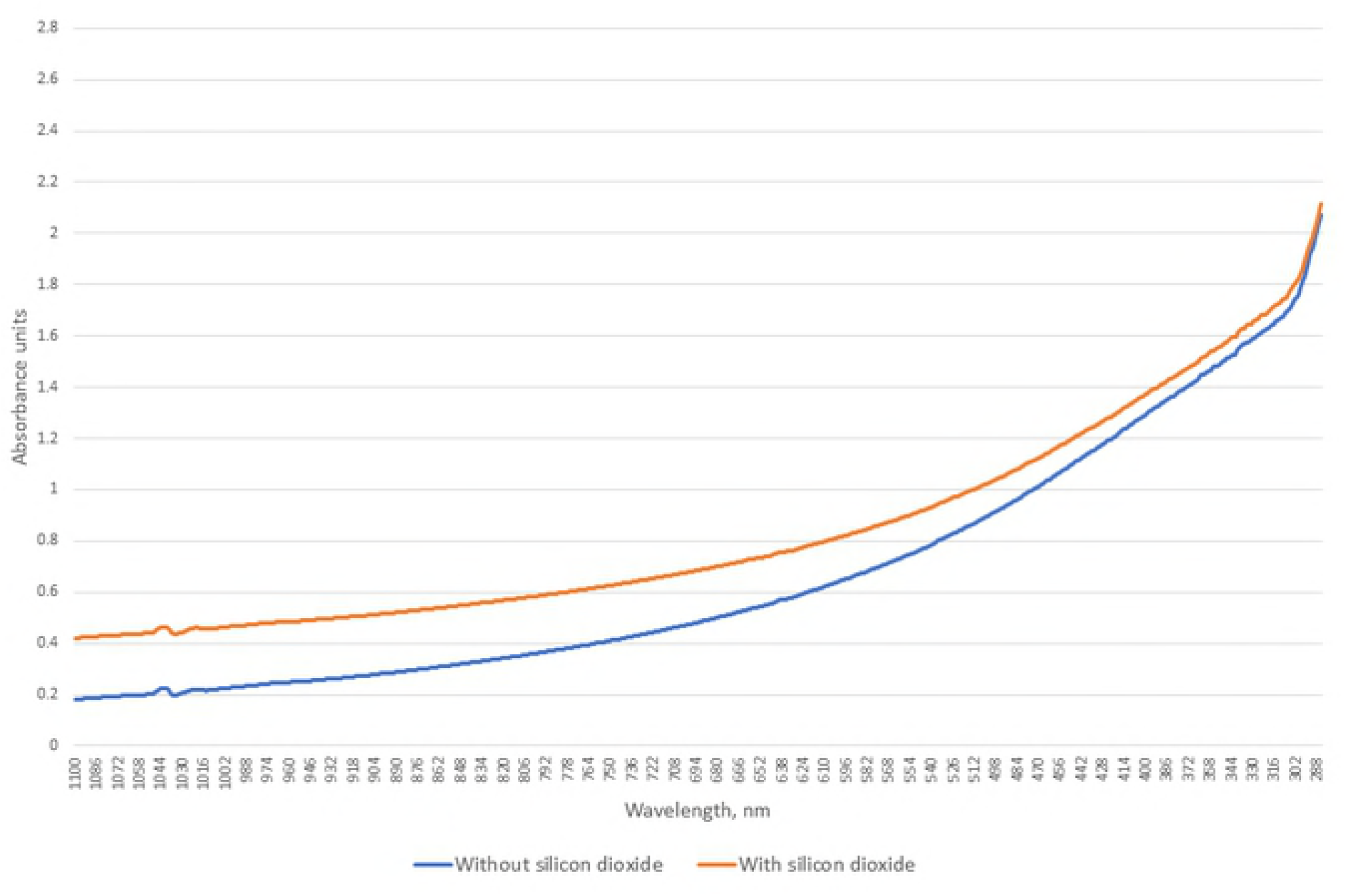
Light absorption of *Klebsiella pneumoniae* with and without silicon dioxide.

**Fig 21.**
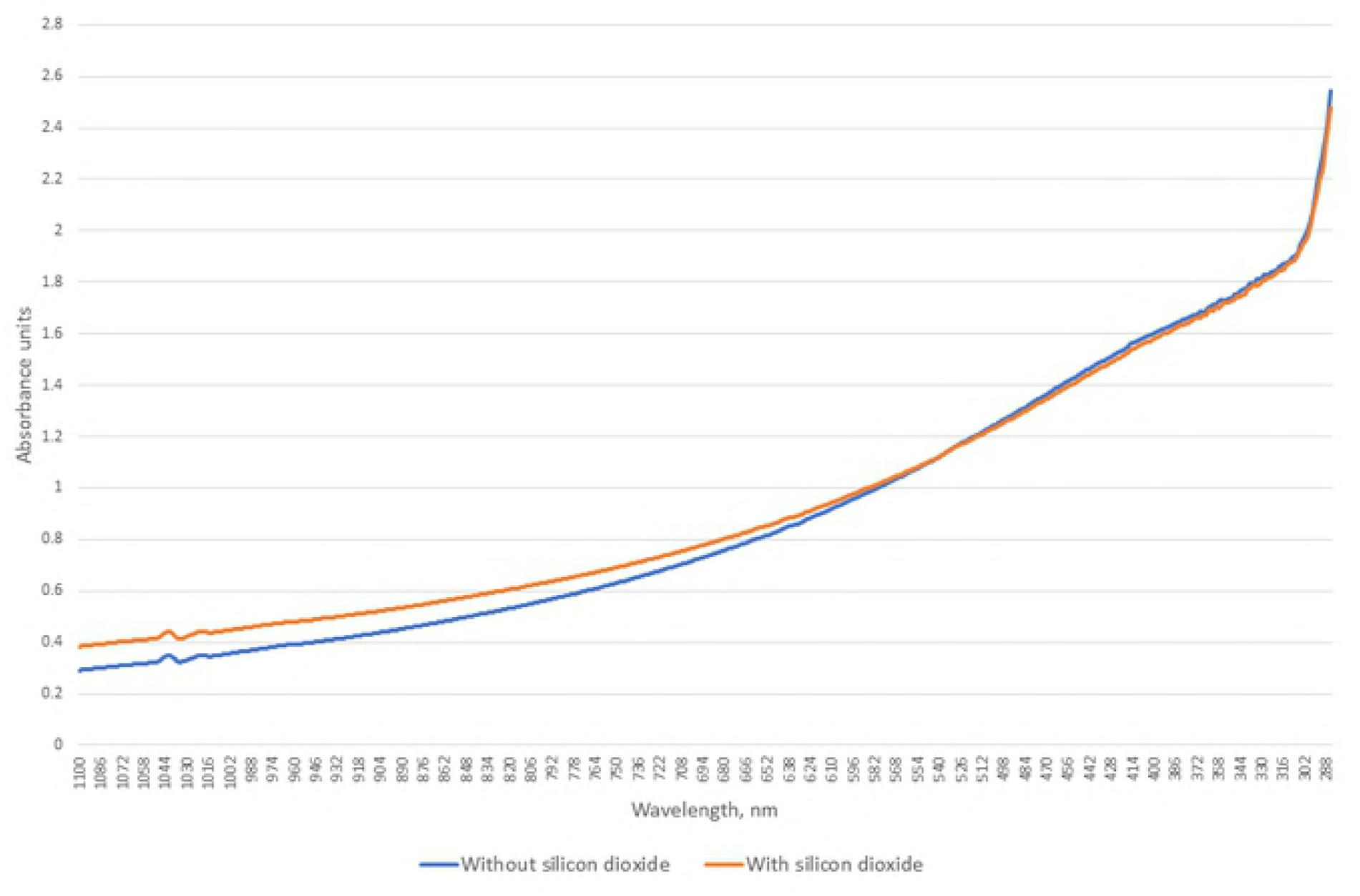
Light absorption of *Moraxella catarrhalis* with and without silicon dioxide.

**Fig 22.**
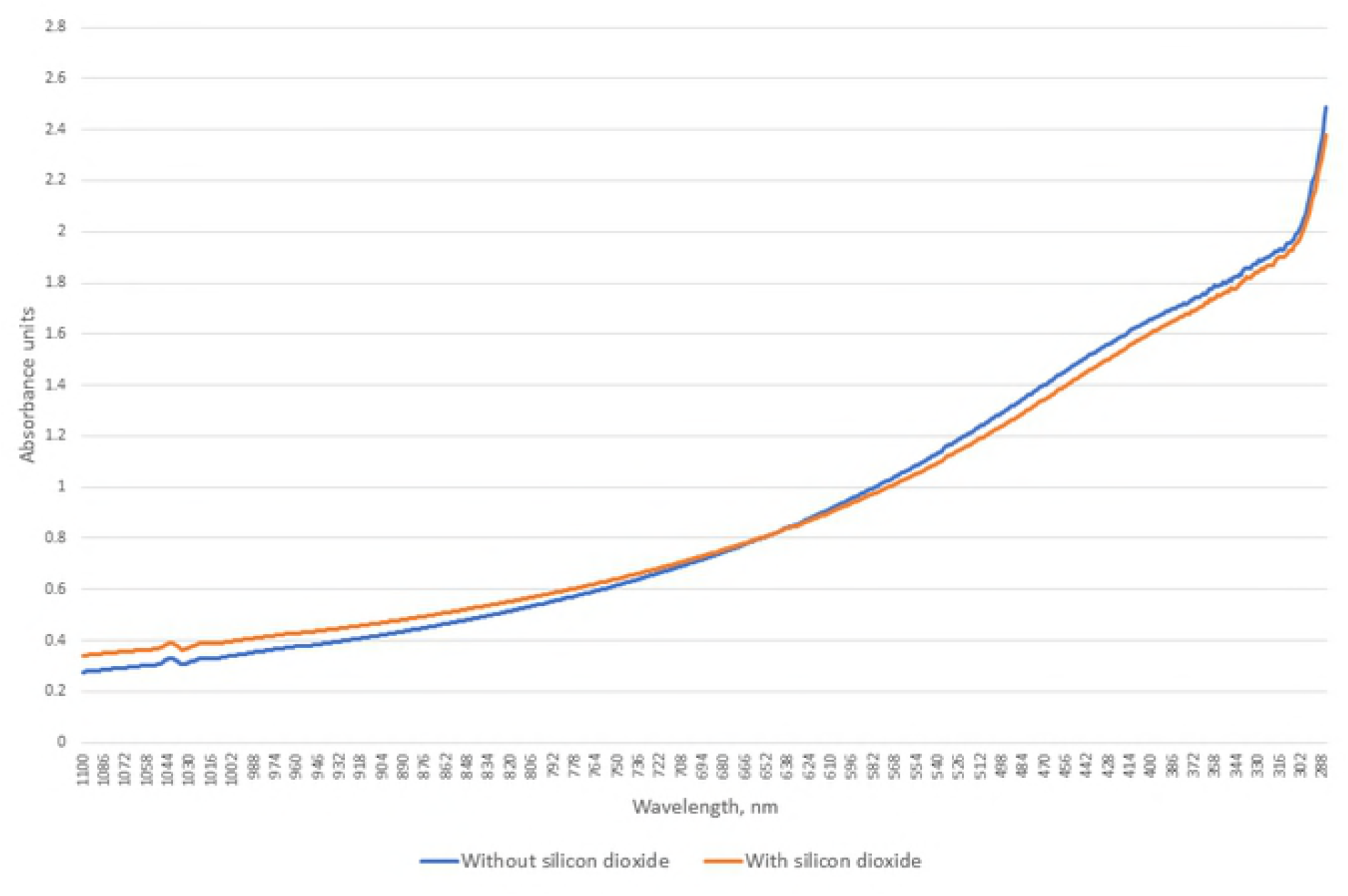
Light absorption of *Escherichia coli* with and without silicon dioxide.

**Table 7.** Effects of silicon dioxide on the light absorption of all examined microorganisms.

| Microorganisms | At what wavelengths does the absorbance increase when adding silicon dioxide |
| --- | --- |
| *Kluyveromyces marxianus* | Lower during the entire measurement |
| *Candida albicans* ATCC | Lower during the entire measurement |
| *Candida glabrata* | Lower during the entire measurement |
| *Candida krusei* | Lower during the entire measurement |
| *Staphylococcus epidermidis* ATCC | 900-1100 |
| *Enterococcus faecalis* ATCC | 690-1100 |
| MSSA ATCC | 605-1100 |
| MRSA ATCC | 430-1100 |
| *Bacillus spizizenii* ATCC | 580-1100 |
| *Proteus mirabilis* ATCC | 610-1100 |
| *Enterobacter aerogenes* ATCC | 550-1100 |
| *Salmonella enteritidis* ATCC | 295-1100 |
| *Pseudomonas aeruginosa* ATCC | 560-1100 |
| *Citrobacter freundii* ATCC | 535-1100 |
| *Klebsiella pneumoniae* ATCC | 285-1100 |
| *Moraxella catarrhalis* ATCC | 540-1100 |
| *Escherichia coli* ATCC | 650-1100 |

## Discussion

### Comparing yeast with bacteria

In this study, we discovered that spectrophotometry could be used to distinguish bacteria form yeast at a wavelength interval of 285-1100 mm by measuring their light absorption. Adding silicon dioxide did not alter the chances to distinguish bacteria from yeast at a wavelength interval of 285-1100 nm, using t-Test: Two-Sample Assuming Unequal Variances. The same could not be said for comparisons made at 285-700 nm. Although visually examining the graph presented by the spectrophotometer, it is possible to distinguish whether or not the examined microorganism is a yeast or a bacteria. The most favourable conditions at which comparisons were made, in an attempt to tell apart a yeast from a bacteria, is at 285-1100 nm without the need to take into account silicon dioxide. It would indicate that a larger wavelength interval is best suited for comparing yeast with bacteria.

### Comparing one bacteria with another

Decreasing the wavelength interval from 285-1100 nm to 285-700 nm improved the odds of distinguishing bacteria apart from one another. At 285-1100 nm without silicon dioxide, only in 52.6% of comparisons it was possible to find a significant difference for the compared bacteria, but at 285-700 nm without silicon dioxide, the results improved to 71.8%, which is a 36.5% improvement. The same thing can be said when silicon dioxide was added. Then it increased from 69.2% to 74.6%, which improved the chances to distinguish bacteria apart by 7.8%. It would indicate that a smaller wavelength interval is best suited for comparing bacteria.

### Comparing one yeast with another

The study showed that the most favourable conditions at which yeast can be compared is at 285-700 nm with silicon dioxide, although silicon dioxide did not provide a substantial light absorption change for yeast. At 285-700 nm without silicon dioxide, in 100% of the comparisons it was possible to distinguish one yeast from another, but adding silicon dioxide the chances dropped to 83.3%, which is a 16.7% drop. Measuring at an interval of 285-1100 nm, in 83.3% of the comparisons it was possible to distinguish one yeast from another, but when silicon dioxide was added the chances drop to 50%. Silicon dioxide gave a 40% drop. The results would indicate that silicon dioxide should not be applied when comparing yeast and a wavelength of 285-700 nm would be a better interval for comparisons than 285-1100 nm.

### Microorganism adhesion with silicon dioxide

Adding silicon dioxide and comparing the changes in the light absorption graphs presented by the spectrophotometer, at long wavelengths the average absorbance of all examined bacteria increased. Although the average absorbance of silicon dioxide is just 0.211 absorbance units. If you would mix a suspension with high absorbance, for example bacteria, with a suspension with low absorbance and then take a sample from the new suspension, you would expect that the light absorption would drop, but for bacteria it was not the case. The increase of light absorption at near infrared wavelengths and for some bacteria at visible light and ultraviolet wavelengths would indicate that bacteria did make complexes with silicon dioxide. For yeast the results are different. The light absorption did drop during the whole measurement for all yeast, with a barely noticeable drop for *Candida glabrata*. It would mean that bacteria did not connect to silicon dioxide or did not form complexes with it that had an altered light absorption. A. Žilēviča and D. Ozoliņš study [13] indicated that yeast do form complexes with silicon dioxide. There for the latter would be a more possible explanations. The results suggest that bacteria have an adherence for silicon dioxide. It would indicate that silicon dioxide is a poor choice of material for the production of prostheses.

Silicon dioxide did not have a significant impact on the average light absorption of *Enterococcus faecalis*, MSSA, *Proteus mirabilis* and *Escherichia coli* (Table 6). It also did not produce any peculiarities in the graphs presented by the spectrophotometer when added to yeast. Which would suggest for a different binding material for the examination of microorganisms with a spectrophotometer.

